# Acclimation of photosynthesis began with a Cu-binding superoxide detoxifying enzyme

**DOI:** 10.1101/2025.07.25.666786

**Authors:** S. Malesinski, A. Vidal-Meireles, E. Giovannetti, M. Chazaux, A. Krieger-Liszkay, C. Boderiou, J. Latil, S. Viola, L.C. Tabares, P. Arnoux, F. Chauvat, P. Dindaeng, C. Maurin, M. Siponen, S. Caffarri, C. Cassier-Chauvat, J. Alric, X. Johnson

**Affiliations:** Aix Marseille Univ, CEA, CNRS, BIAM, UMR7265; Saint-Paul-Lez-Durance, 13115, France; Institut de Biologie Intégrative de la Cellule (I2BC), CEA, CNRS, Université Paris-Saclay; Gif-sur-Yvette, 91190, France; Aix Marseille Univ, CEA, CNRS, BIAM, UMR7265, Luminy Plant Genetics and Biophysics Team; Marseille, 13009, France

## Abstract

Plant acclimation is a growing scientific concept, at molecular, cellular and global scales. All photosynthetic organisms that created an oxic atmosphere on earth possess a gene of unknown function “*Acclimation of Photosynthesis to the Environment 1*”. Here we show that *APE1* encodes a thylakoid-bound protein with a unique motif that binds copper and detoxifies the superoxide anion radical, O_2_^•−^. Maturation of the recombinant APE1 protein from *Chlamydomonas reinhardtii* requires formation of cysteine disulfide bonds after copper binding or *via* a high affinity interaction with a copper chaperone (Plastid Copper Chaperone 1) that boosts its scavenging capacity for O_2_^•−^. APE1 co-occurs in evolution with Photosystem II oxygen evolving proteins and it is the archaic O_2_^•−^ detoxifying enzyme for acclimating photosynthesis to an oxygenic environment.

## Main text

Over time, vegetation responds to environmental changes, such as light, temperature and CO_2_ availability by adjustments to its physiology. This is referred to as plant acclimation, and it is a growing concept in science, in various fields and from the molecular scale (*1*) to the global level (*2*). Acclimation of photosynthesis is now implemented in land surface models (LSM) with an aim at deriving a more general response of vegetation across plant functional types, thereby simplifying the parametrization of the models (*3–5*). The trait “*Acclimation of Photosynthesis to the Environment 1*” (*APE1*) was identified in genetic screens in *Arabidopsis* and *Chlamydomonas* (*6,7*) where mutants could not maintain Photosystem II quantum yield when exposed to a higher light intensity. Various proteomics studies have found APE1 linked to the D2 module during PSII biogenesis (*8–10*), but its Domain of Unknown Function 2854 has remained undefined. Nevertheless, the strict conservation of APE1 in all oxygenic phototrophs (*6*) and its involvement in light acclimation suggest an ancestral role in PSII maintenance and photoprotection.

Oxygenic photosynthesis produces dioxygen (O_2_) from water splitting at the level of Photosystem II. Along the photosynthetic electron transport chain, strongly reducing reaction intermediates can reduce O_2_ into the superoxide anion radical (O_2_•−), at level of Photosystem I (*11*), Photosystem II (*12,13*) or cytochrome *b*_6_*f* complex (*14*). O_2_•− is a reactive oxygen species (ROS) that is toxic for living cells. Superoxide dismutases (SOD) detoxify O_2_^•−^, working at diffusion-limited rates to produce O_2_ and H_2_O_2_ from 2 molecules of O_2_^•−^ (*15*). The soluble Fe- SOD isoforms in the stroma of chloroplasts, conserved across species, are the first step of the “water-water” cycle to detoxify the O_2_^•−^ produced by PSI (*16*). However, detoxification at other O_2_^•−^ production sites along the thylakoid membrane remains virtually unknown. Enzymatic O2^•−^ scavenging *via* the protein Antioxidant 1 (ATX1) and related homologues (*17–19*) has also been reported in yeast, plant cytosol and animals.

In this work, we show that APE1 has its origins at the beginnings of oxygenic photosynthesis. To find the function of the DUF2854, we produced the recombinant versions of the APE1 orthologues from three very different photosynthetic organisms, the cyanobacterium *Synechocystis* PCC 6803 (slr0575, *SyAPE1*), the green algae *Chlamydomonas reinhardtii* (Cre16g665250, *CrAPE1*), and the angiosperm *Arabidopsis thaliana* (At5g38660, *AtAPE1*). We showed that APE1 binds copper *in vitro. Via* site-directed mutagenesis of the active and regulatory sites of APE1 from *Chlamydomonas* (*Cr*APE1), we reveal its complex dual maturation pathway. We show the Plastid Cu Chaperone 1 (PCC1) transfers copper to *Cr*APE1, and provide evidence that the two proteins act as plastid antioxidants, cooperatively participating in acclimation of the photosynthetic apparatus to high light *via* their O_2_^•−^ scavenging activity. Our results suggest that APE1 would have played an important role in the early steps of the evolution of the oxygenic photosynthetic apparatus, protecting it from its potentially harmful by-product.

### *APE1* has an evolutionary link to PSII and is found in all oxygenic phototrophs

*APE1* is a GreenCut gene, conserved only in oxygenic photosynthetic organisms, from cyanobacteria to plastid-containing eukaryotes (*20,21*). Comparative genomic analyses also identified *APE1* as part of the cyanobacterial core genome, which consists of only 63 genes shared amongst all oxygenic photosynthetic organisms (*22*). We extended this comparison using 77 cyanobacterial genomes, including those that have lost PSII and the genes required for carbon fixation (*23,24*). Through this cross-analysis, *APE1* was restricted to PSII-containing cyanobacteria in a group of nine genes with a similar co-occurrence that included genes coding for PSII core subunits and the O_2_ evolving complex (table S1). This early appearance and conservation of *APE1* in photosynthetic organisms, suggests that APE1 played a pivotal role in the emergence of O_2_ evolution.

### The membrane-bound APE1 protein has a domain of unknown function

In all species, APE1 has 2 transmembrane helices (TMH). In eukaryotes it has a chloroplast transit peptide (fig.1A). The soluble stromal 15-17 kDa domain annotated as unknown function (DUF2854) has a conserved motif DVTR(Y/H)RYGDE(A/Q)HL(D/E) (inset, fig.1A). It is rich in charged amino acids (H, D and E), reminiscent of a metal-binding site or an enzymatically active site. D152, Y158, and H163 (*Cr*APE1 numbering) are conserved in all the orthologues. The majority of prokaryotes (*Synechocystis*) have no cysteines (C), while unicellular eukaryotes (*Chlamydomonas*) have up to 2 C and more complex, multicellular organisms (*Arabidopsis*) up to 3 C (fig. S1). Like for many other proteins an increase in C number in APE1over the course of evolution suggests additional roles related to function, regulation, oligomerization or interactions (*25*). The cysteine C128 is conserved throughout the green lineage and in some diatoms (SF1).

**Figure 1.**
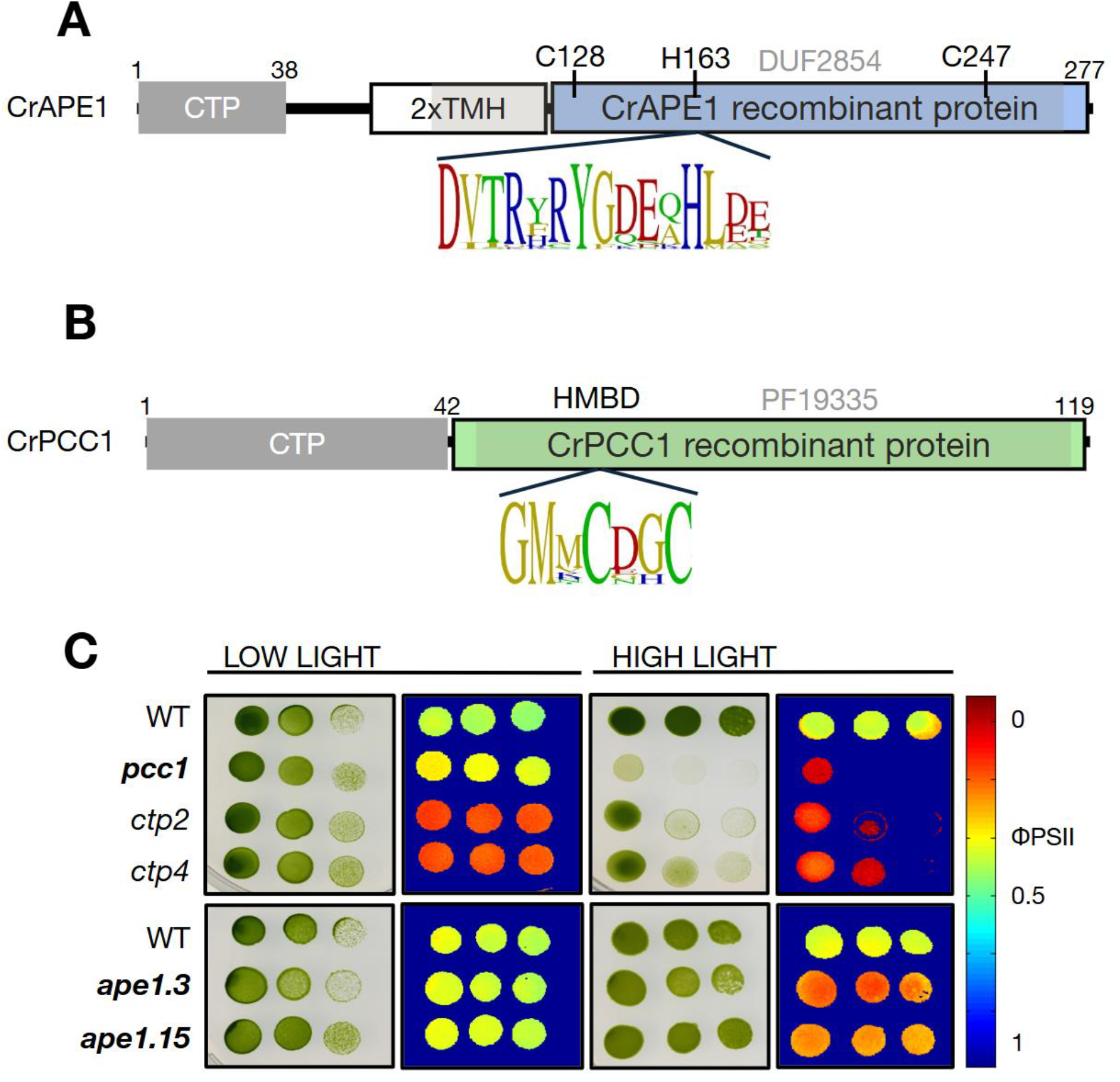
Proteins *Cr*APE1 and PCC1 are Required for Light Acclimation of Photosynthesis. **A**. APE1 protein model (*Chlamydomonas reinhardtii*) contains chloroplast target peptide (CTP), two thylakoid transmembrane helices (TMH) and the DUF2854 (grey shading) notably containing cysteines at positions 128 and 247 and the consensus sequence around the histidine 163. B. Plastid Copper Chaperone 1 (PCC1) protein model (*Chlamydomonas reinhardtii*) shows the heavy metal binding domain HMBD (Pfam PF19335; gray shading) and its consensus sequence in PCC1 orthologues. C. Growth tests and imaging of photosynthetic activity under low light and high light phototrophic conditions. Mutants of *pcc1* (bold) and copper transporters *ctp2* and *ctp4* are compared against their corresponding WT strains and *ape1* mutants (bold).

### *Plastid Copper Chaperone 1* (*PCC1*) is co-expressed with *CrAPE1*

To provide context for *APE1* function, data-mining of the top 250 genes co-expressed with *Cre16*.*g665250* (obtained from PhytoMine against *C. reinhardtii* genome V5.6: https://phytozome-next.jgi.doe.gov/phytomine/begin.do) showed high representation from green cut proteins targeted to the plastid (*20*). These factors could be grouped into functional clusters involved in assembly or repair of photosynthetic complexes, thylakoid remodeling, alternative electron pathways, carbon metabolism, redox and antioxidant regulation, pigment biosynthesis, and cofactor/chaperone activity (fig. S2). These same functional groups could be found in *APE1* co-expression data from *Synechocystis sp. PCC 6803* (ALCOdbCyano) and *Arabidopsis* (phytozome), establishing the case for a similar function across very diverse photosynthetic organisms (fig. S2). Among the genes co-expressed with *CrAPE1* we found Cre05.g248600, annotated as *Plastid Copper Chaperone 1* (*PCC1*)(*26*). This gene model contains a predicted plastid target sequence and the Pfam Heavy Metal Binding Domain (HMBD, PF19335), a conserved Cu-binding motif GMXCXXC (Fig.1B). Orthologues of *Cr*PCC1 may only be retained in the *Chlamydomonales* (*Chlamydomonas* and *Volvox*) but other *Chlorophytes* and diverse bacteria have a protein with significant sequence similarity (fig. S3). PCC1, as well as its closely related proteins, showed high structural homology to the ATX1-like Cu chaperones, such as CopZ, ATX1, and Domain 1 of the Copper Chaperone to SOD (CCS), containing the typical ferredoxin-like fold (βαββαβ) (fig. S4), the HMBD and a number of conserved lysines (K).

### *Cr*APE1 and PCC1 maintain high PSII quantum yield in high light

We measured the acclimation from low light to high light (fig. 1C) of *Chlamydomonas* mutant strains *ape1* and *pcc1*, and of *ctp2* and *ctp4* that are devoid of P-type ATPase Cu transporters localized to the chloroplast envelope and the thylakoid membrane, respectively. All strains have insertions in exons of the genes (fig. S5). The *ctp2* and *ctp4* mutants have reduced growth and low PSII quantum yield (Φ_II_) even under low light, resulting from low amounts of the Cu-containing electron carrier plastocyanin (fig. 1C and fig. S6). Interestingly, the loss of PCC1 does not affect plastocyanin levels (fig. S6). The *pcc1* and *ape1* mutants are less affected in low light, but show impaired acclimation to high light, with decreased Φ_II_ in both, and slower growth in *pcc1*. Hence, while CTP2 and CTP4 are required for plastocyanin accumulation, PCC1 is likely involved in a different pathway in the chloroplast, such as Cu sequestration, Cu transport or antioxidant activity, as reported for ATX1 (*19*). The co-expression and similar acclimation response to high light suggested that PCC1 and APE1 might be involved in the same response mechanism.

### APE1 is a copper-binding protein

We expressed the soluble region of the domain of unknown function, DUF2854, of *Synechocystis* PCC6803 (*Sy*), *Chlamydomonas* (*Cr*) and *Arabidopsis* (*At*) as recombinant proteins in *E*.*coli* (fig S1 and S7 and table S2). We tested APE1’s interaction with divalent cations using nano differential scanning fluorimetry. Protein fluorescence was quenched at 330 nm and shifted to 350 nm upon CuCl_2_ addition. This was observed as an increase in the 350/330 nm ratio for *Cr*APE1 (fig. 2A), *At*APE1 and *Sy*APE1(fig. S9 A-F), and lowered the melting temperature (T_m_) (fig. 2B and fig. S9 G-I). Of all the divalent cations tested, only Zn had the same effect as Cu on the T_m_, possibly suggesting a promiscuous binding site, a second site, or a non-specific interaction. In all other instances, the changes in protein fluorescence were specific to Cu and dose-dependent, suggesting that protein conformational changes were induced by Cu binding. We then measured Cu affinity at lower protein concentrations and found high affinities 128 < *K*_D_ < 580 nM (fig. S10). Notably, the addition of reductant (TCEP) inhibited Cu-binding for *Cr*APE1 (see inset fig. S8) either due to the reduction of Cu(II) to Cu(I) or to the reduction of cysteine disulfide bonds (S-S) to free thiols (−SH). The most common residues binding Cu are H and C (MetalPDB: https://metalpdb.cerm.unifi.it/), thus we used EPR and site-directed mutagenesis in *Cr*APE1 to identify the Cu coordination.

**Figure 2.**
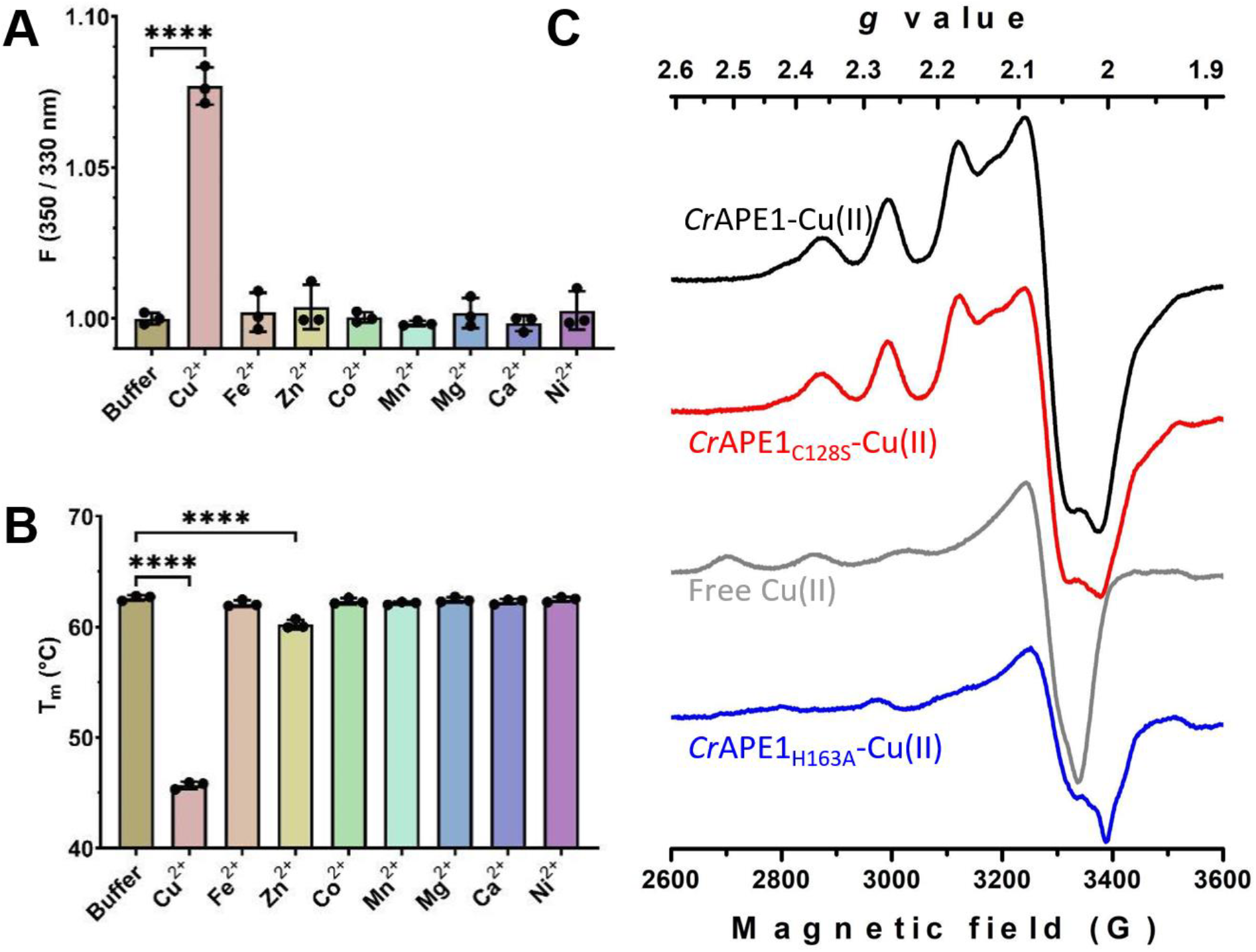
*Cr*APE1 recombinant protein specifically interacts with Cu(II) via H163. **A**. Cu(II) significantly changes protein conformation, measured as an increase of the fluorescence ratio of aromatic residues (350/330 nm). B. Cu decreases the melting temperature of *Cr*APE1, so does Zn to a lesser extent. The statistical relevance is shown as pairwise comparisons (*p*≤0.0001 ****). C. X-field EPR spectra of *Cr*APE1WT-Cu(II) and *Cr*APE1_C128S_-Cu(II) mutant are different from free Cu(II) and the *Cr*APE1_H163A_-Cu(II) mutant showing that H163 binds copper.

### Cu binds to the conserved Histidine 163

The Cu(II) EPR spectrum of *Cr*APE1_WT_ recombinant protein showed an anisotropic shape (thick black line in Fig. 2C), different from free copper (grey). To define the binding site, we analyzed the Cu(II) EPR spectra of *Cr*APE1 where the conserved histidine was substituted with alanine (*Cr*APE1_H163A_) and in the mutant of the most conserved of the two cysteines (*Cr*APE1_C128S_). The spectrum of the *Cr*APE1_C128S_ mutant (red) closely resembled that of the wild type, while the spectrum of the *Cr*APE1_H163A_ mutant (blue) was significantly different. However, the H163A spectrum was still distinct from that of Cu(II) in solution (grey). These results show that the only histidine, His 163, binds Cu(II) but C128 does not. Specific binding of Cu(II) to H163 was confirmed by mass spectrometry, (table S2 and fig. S12).

### Regulation of *Cr*APE1 through disulfide bonds

In size-exclusion chromatography (SEC), *Cr*APE1-Cu eluted earlier than the apoprotein (fig. 3A), supporting the change in protein conformation suggested from protein fluorescence (see above). *Cr*APE1-Cu was predominantly a monomer as confirmed by multi-angle light scattering (MALS) (fig. 3C), with < 6% of the molecules forming dimers and other oligomers (table S3). Any changes to mobility or oligomerization were suppressed when the protein was pretreated with TCEP, further substantiating the absence of fluorescence quenching in this condition (as in fig. S8). In order to discriminate between the possible redox effect of TCEP on Cu(II) / Cu(I) or on the two cysteines C128 and C247, we compared WT to C128S mutant (fig. 3B). In the *Cr*APE1_C128S_ mutant, the monomer did not shift upon Cu addition and no differences were observed in the TCEP-treated sample. This suggested to us that intramolecular disulfide bond formation (S-S) occurred in APE1_WT_. However, *Cr*APE1_C128S_-Cu formed a larger molecular complex, (fig. 3B and ST3), identified as a protein dimer by MALS (fig. 3D). This suggested that C247, normally engaged in an intramolecular disulfide bond with C128 in the WT, became more available for an intermolecular disulfide bridge and stabilized the protein dimer in *Cr*APE1_C128S_-Cu. TCEP fully dissociated this dimer (seen in fig. 3B), further substantiating the role of C247 in the interaction between monomers.

**Figure 3.**
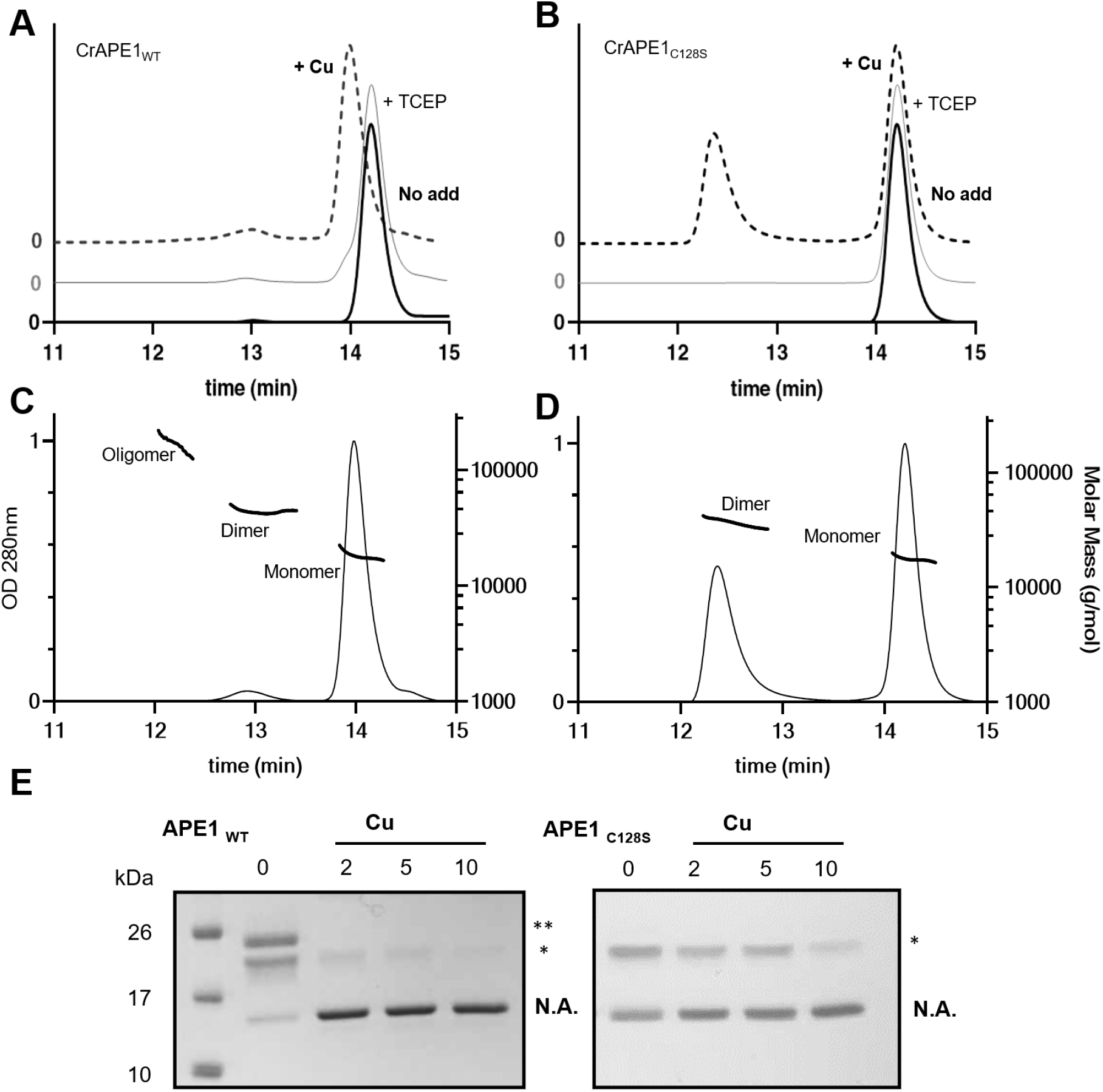
The cysteines of *Cr*APE1 control monomerization through Cu-binding to His163. Analysis of *Cr*APE1 oligomerisation: size exclusion chromatography (SEC) profile of **A**. *Cr*APE1_WT_ and B. *Cr*APE1_C128S_ incubated without additive (black), with CuCl_2_ addition in a ratio of 1/5 (dotted) or with 1mM TCEP followed by CuCl_2_ addition in a ratio of 1/5 (grey). Molecular mass was estimated by multi-angle light scattering (MALS) for C. *Cr*APE1_WT_ and D. *Cr*APE1_C128S_. *Cr*APE1_WT_ and *Cr*APE1_C128S_ alkylation with mPEG-maleimide-2000. E. The number of cysteine thiols was tested by PEGylation and indicated by asterisks (*). N.A. refers to non-alkylated protein.

To confirm the redox state of the cysteine residues in *Cr*APE1, we used an SDS-PAGE mobility-shift assay by alkylation with mPEG. In non-treated *Cr*APE1_WT_, containing two C, three bands were found after alkylation (fig. 3E): *Cr*APE1 alkylated with two PEG tags (**), with one PEG- alkylation (*), and non-alkylated (N.A.) protein. After addition of Cu, *Cr*APE1_WT_ was non alkylated, showing that cysteines were oxidized, forming an intramolecular disulfide bridge (S-S). Expectedly, *Cr*APE1_C128S_ formed only two bands, non-alkylated and alkylated, as it contains only one C. The alkylation of the sole C was maintained upon addition of Cu. Altogether, these results indicate that Cu-binding to *Cr*APE1_WT_ stabilizes an intramolecular disulfide bridge between C128 and C247. In the absence of C128, a disulfide bond between C247 residues stabilizes the homodimer.

### *Cr*APE1 has high affinity for Plastid Copper Chaperone 1 (PCC1)

Addition of free Cu to the growth medium during protein expression neither increased *Cr*APE1 yield nor enhanced Cu loading in the purified protein, suggesting the requirement for a Cu chaperone. We thus produced recombinant PCC1 (in the presence of Cu). We identified Cu-PCC1 as a 16 kDa homodimer, with a small fraction of 8 kDa monomer (fig. S4D,E). Metal analysis by MP-AES showed it contained 0.59 Cu per PCC1, making 1 Cu atom per PCC1 dimer (table S4). This resembles ATX1 copper chaperones, where Cu is coordinated by 4 cysteine residues, 2 from each monomer (*27*). Microscale Thermophoresis (MST) confirmed an interaction between CuPCC1 and *Cr*APE1_WT_ with a dissociation constant of *K*_D_ = 40 nM (fig. 4A), virtually unchanged in *Cr*APE1_C128S_ (*K*_D_ = 38.6 nM). Similarly to ATX1, PCC1 likely interacts with APE1 through electrostatic interactions via a conserved set of 6 lysines (*19*) (fig. S3).

**Figure 4.**
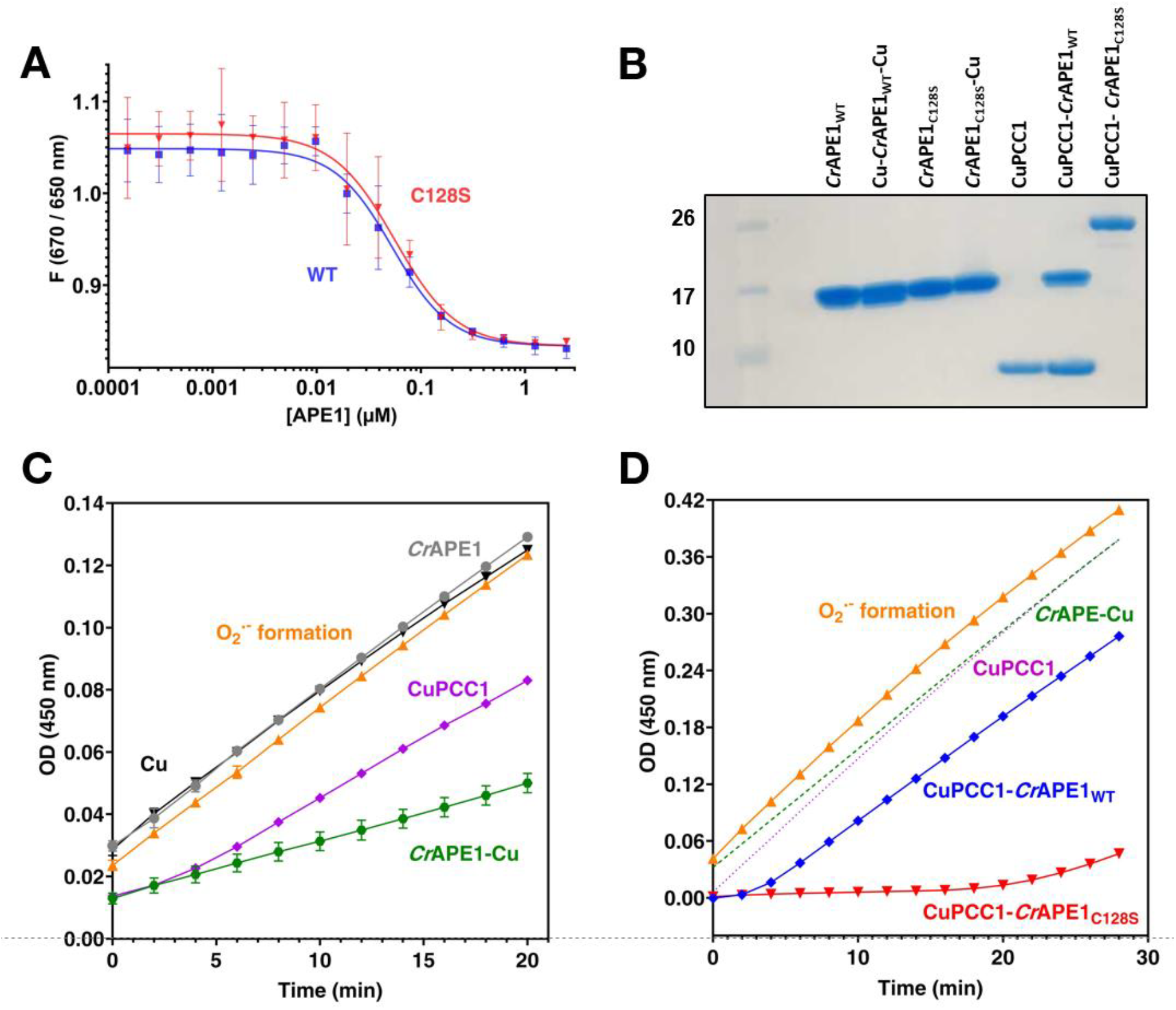
APE1 and PCC1 interact and detoxify superoxide. **A**. PCC1 binds *Cr*APE1 (*K*_D_ = 40 nM) and this is independent from the C128 residue: *Cr*APE1_WT_ (blue), *Cr*APE1_C128S_ (red). B. Denaturing SDS-PAGE separation of *Cr*APE1_WT_, *Cr*APE1_C128S_ and PCC1 alone and in complex with Cu and co-purified. C. *Cr*APE1 incubated with Cu (green) at 50% O_2_^•−^ detoxification rates has sustained activity; PCC1 (purple) at 50% O_2_^•−^ detoxification has scavenging activity, observed by the biphasic slope. Negative controls are *Cr*APE1 without Cu (gray) and Cu alone (black). D. O_2_^•−^ scavenging by CuPCC1-*Cr*APE1 wildtype (blue) and C128S mutant (red) complexes. The initial rate of superoxide detoxification exceeds the rate of production, read as 100% detoxification, then slows. Additional controls *Cr*APE1-Cu (green) and PCC1 (purple) are shown for comparison. All proteins are at concentrations of 75 µM. For biological triplicates, see supplementary data set 2. O_2_^•−^ was generated by xanthine/xanthine- oxidase and reported by the dye WST-1 for which absorbance was followed at 450 nm; the orange trace in C. shows formazan formation from O_2_^•−^ production.

### *Cr*PCC1 acts as a Cu metallo-chaperone for *Cr*APE1

Such high affinity between these two proteins suggested that PCC1 was involved in delivering copper to *Cr*APE1. To test this, we expressed the two proteins in two separate *E. coli* cultures, induced in the presence of Cu for PCC1 or in absence for *Cr*APE1. We co-purified *Cr*APE1_WT_ and CuPCC1 by lysing *E. coli* cultures together then proteins were dialyzed and separated by cation exchange chromatography (fig. 4B; fig. S13). MP-AES detected Cu in all analyzed fractions (ST 4). The pre-separated CuPCC1-*Cr*APE1_WT_ complex contained 1 Cu per complex. After separation, all *Cr*APE1_WT_ fractions contained Cu and some fractions up to 0.84 Cu per *Cr*APE1. This demonstrates that CuPCC1 can transfer Cu to *Cr*APE1 *via* a mechanism that has not been previously described.

### Stabilization of the transient CuPCC1-*Cr*APE1 complex formed upon Cu transfer

We co-purified *Cr*APE1_C128S_ with CuPCC1 in an identical manner. When the wildtype and mutant complexes were separated by SDS-PAGE with addition of DTT, the CuPCC1-*Cr*APE1_WT_ complex dissociated as 8 kDa and 17 kDa bands but CuPCC1-CrAPE1_C128S_ did not, running as a single band at 25 kDa (fig. 4B). The complex formed by CuPCC1-*Cr*APE1_C128S_ was thus resistant to dissociation by reducing or denaturing agents. This suggested that co-purified CuPCC1 and *Cr*APE1_C128S_ were trapped in the maturation process providing us with a snapshot of an intermediate. The most likely explanation is that CuPCC1-*Cr*APE1_C128S_ formed a Cu-dithiolate complex through the C247 residue. When in interaction with APE1_WT_, the complex was only transient because it dissociated through competition with H163 for the Cu and with the thiol group of Cys128 for the protonation and release of PCC1 (a model is presented in fig. 5).

**Figure 5.**
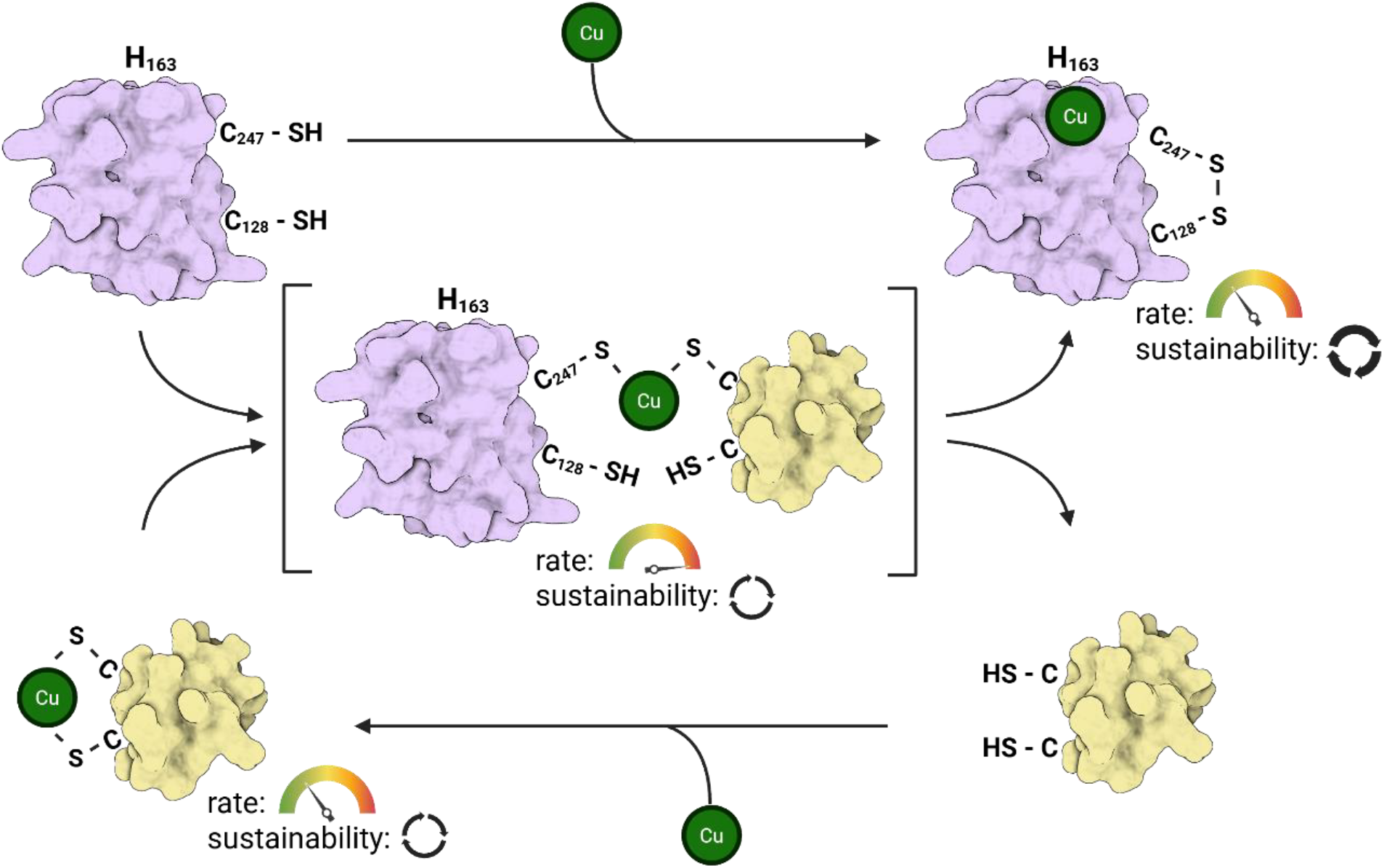
Model for APE1 copper incorporation and enzyme activity. Copper uptake by APE1 may vary, from species to species, as for CuZn SOD1. From cyanobacteria to green algae and higher plants, nascent APE1 spontaneously binds Cu. Cu binding to APE1 induces protein conformational changes and a disulfide bond is formed between C128 and C247 in *Cr*APE1, this provides APE1 with a low but sustained O_2_^•−^ detoxification activity. *Cr*APE1 is matured with the PCC1 Cu-dithiolate complex, homolog to ATX1 or domain 1 from human CCS. Upon interaction, the C247 thiol attacks PCC1 Cu-thiolate, forming a PCC1-APE1 Cu-dithiolate heterodimeric complex. Then C128 thiol attacks the second PCC1 Cu-thiolate, thereby transferring Cu to the His-binding site in the matured APE1 and forming an intramolecular disulfide bond. This dissociates the CuPCC1-*Cr*APE1 complex and makes PCC1 available for subsequent Cu binding. PCC1-Cu is a low activity O_2_^•−^ scavenger. The C128S substitution prevents the second nucleophilic attack of the Cu-thiolate and stabilizes the CuPCC1-*Cr*APE1 heterodimer. This complex is a high activity O_2_^•−^ scavenger. Created in AlphaFold3 and BioRender.com

### *Cr*APE1-Cu and CuPCC1 have superoxide detoxification activity

Based on the uncanny similarities to the CCS1 maturation pathway of CuZnSOD, we tested the reactivity of the recombinant proteins with O_2_^•−^. We generated O_2_^•−^ by xanthine/xanthine oxidase and used the dye WST-1 formazan to report O_2_^•−^ formation as an absorbance increase at 450 nm. *Cr*APE1-Cu (122 µM) inhibited 50% of O_2_^•−^ production, steadily over the 20-minute assay, while apoAPE1 (150 µM) or free copper showed no reaction with O_2_^•−^ (fig. 4C). CuPCC1 inhibited 50% of O_2_^•−^ production with less protein (50 µM) but O_2_^•−^ detoxification was not sustained and resembled the rates of scavenging of O_2_^•−^ previously described for ATX1 (*18,19,28*). The enzymatic activity for both CuPCC1 and CrAPE1-Cu was thus 2-3 orders of magnitude lower than the CuZnSOD (*15*). We then tested CuPCC1-*Cr*APE1_WT_ (blue) and again found strongly biphasic kinetics but at higher production rates of O_2_^•−^ (fig. 4D). CuPCC1-*Cr*APE1_WT_ had an initial phase of full O_2_^•−^ detoxification over 3 minutes, followed by a slower phase of sustained activity (blue) that correlated with the slope of *Cr*APE1_WT_ (green). This initial phase of full O_2_^•−^ detoxification was extended to 20 min in CuPCC1-*Cr*APE1_C128S_ (red). The extent of the fast detoxification phase (Fig. 4D) correlated to the amount of CuPCC1-*Cr*APE1 stable complex (Fig. 4B), and not to the activity of CuPCC1 (purple). This suggests that PCC1 has a dual function both in maturation and in transiently boosting O_2_^•−^ detoxification of *Cr*APE1 (see Fig. 5).

## Discussion

Here we identified the function of the first molecular factor involved in acclimation of photosynthesis to the environment, APE1, a chloroplast thylakoid copper-binding enzyme detoxifying the superoxide anion radical O_2_^•−^at the vicinity of photosynthetic complexes. The CuPCC1-*Cr*APE1 interaction results in a transient but augmented activity that may be relevant under stress conditions. We suppose that APE1 alone, found in nascent PSII assembly intermediates, would have sufficient activity to protect the cyt *b*559 from O_2_^•−^ production (*12,29*), when in complex with D2 (*8,9*). While the higher O_2_^•−^ scavenging capacity through PCC1-APE1 interaction could be important for protection of mature PSII when the non-heme Fe is devoid of bicarbonate (*13,28*). There is also strong evidence of functional activation of APE1 with other proteins and pathways, with its inclusion in autophagy stromules and plastoglobules under light stress (*30,31*). Given these interactions, APE1’s function in O_2_^•−^ detoxification reveals a pivotal role in chloroplast acclimation to stress.

ROS detoxification enzymes enabled life on Earth to transition from an anoxic to an oxic environment and living cells have expanded their repertoire of enzymes. Superoxide dismutases and reductases (SOR) use various elements Ni, Fe, Mn or Cu/Zn as cofactors for electron transfer (*15*). Although these metals seem equally effective for catalysis, their availability as micronutrients changed dramatically during earth history (*32*). During the Great Oxidation Event, oxygenic photosynthesis precipitated Fe(II) into Fe(III) and solubilized copper sulfides, making Cu(II) more available than Fe(III) (*32,33*). It seems consistent that APE1, which appeared at this time in evolution, selected Cu(II) as a protein cofactor (*33*). Later in evolution in a green alga, it recruited the Cu chaperone PCC1, an atypical, chloroplastic, ATX-like protein whose transcription decreases in response to Cu deficiency (*26*).

APE1 dates back to oxygen production as far as we can trace it. Independent genetic screens have identified it as the locus *Acclimation of Photosynthesis to the Environment 1* (*APE1*) (*6,7*). In high light, when photosynthesis is limited by CO_2_ availability, the resilience of Photosystem II depends on APE1 (*6,7*). APE1’s substrate, O_2_^•−^, is not only a harmful byproduct of photosynthesis, toxic for living cells, it is also used as signal molecule for biological acclimation (*34,35*). In this context, APE1 stands as the first example of a regulatory control point of O_2_^•−^ at the intersection of photosynthesis, light and CO_2_ levels, fully justifying its identification as *Acclimation of Photosynthesis to the Environment 1*.

## General

We acknowledge the technical help from Tiffanie Barre, Patricia Henri, Geraldine Brandelet and Clara Ansellem. We would like to thank Sebastien Thomine (I2BC) for his help with MP-AES. We warmly thank Bernard Genty for fruitful discussions.

## Funding

Agence National de la Recherche grant « ChloroPaths » ANR-14-CE05-0041-01 (XJ)

Agence National de la Recherche grant « RevelOrg » ANR-20-CE20-0006 (SC, CCC, XJ)

Labex Saclay Plant Sciences (SPS) grant ANR-17-EUR-0007 (AKL)

I2BC Biophysics Platform FRISBI IBiSA, grant ANR-10-INSB-05 (LT and AKL) European Union Horizon 2020 grant “CAPITALISE” 862201 (JA and XJ)

PEPR FairCarboN National Research Agency - France 2030 grant « GREENSCALE » ANR-23-PEXF-0003 (XJ, JA)

## Author contributions

Conceptualization: SM, AVM, MC, CB, JL, SV, PA, FC, SC, CCC, JA, XJ

Methodology: SM, AVM, EG, MC, AKL, CB, JL, SV, LT, PA, PD, CM, CCC, MS, XJ

Investigation: SM, AVM, EG, MC, AKL, CB, JL, SV, LT, PA, PD, CM, CCC, MS, JA, XJ

Funding acquisition: XJ, JA, AKL, CCC, SC

Project administration: XJ

Supervision: XJ, SM, JA, CCC, SC

Writing – original draft: XJ, JA, AVM, SM

Writing – review & editing: SV, PA, CCC, FC, SC, AKL, LT, JA, XJ

## Competing interests

Authors declare that they have no competing interests.

## Data and materials availability

All data, and materials used in the analysis are available to any researcher for purposes of reproducing or extending the analysis.

## Supplementary Materials

Materials and Methods

Figs. S1 to S13

Tables S1 to S5

References

(22-24 and 36-43)

Data S1 to S2

## Notes

### Competing Interest Statement

The authors have declared no competing interest.

