## Supplementary material for "Acclimation of photosynthesis began with a Cu-binding superoxide detoxifying enzyme": M&M and Supplemental figures and tables

**Affiliations:**

**The PDF file includes:**

Materials and Methods
Supplementary Figures S1 to S13
Tables S1 to S5
References in Supplementary files (also included in main references)

**Other Supplementary Materials for this manuscript include the following:**

Data S1. *APE1* Co-expression data in 3 species.
Data S2. Superoxide detoxification assays

### Materials and Methods

Algal cell culture.

The wild-type strain (T222 $mt^+$) used for generating the *ape1* knock-out mutants is a progeny of 137c backcrosses (*36*); CC-5325 background wild-type strain and *pcc1* (*LMJ.RY0402.059810*), *ctp2* (*LMJ.RY0402.149111*), and *ctp4* (*LMJ.RY0402.047261*) mutants were obtained from the CLiP mutant collection background (*37*). Maintained cultures were always cultivated at 20 µmol photons·$m^{-2}$·$s^{-1}$ in TAP + 2 % agar media plates. For experimental setups, liquid cell cultures were grown in Tris-Acetate-Phosphate (TAP) medium at 25°C under ambient air at continuous 25 µmol photons·$m^{-2}$·$s^{-1}$ illumination in incubation shakers (photomixotrophic conditions) for 3 days, and the shifted to photoautotrophic conditions by centrifugation and resuspension in minimal (MIN) media under ambient air before being plated for spot tests, photos were taken at 7 days. Standard recipes for media preparation were used as in (*38*).

Generation of *C. reinhardtii* CRISPR/Cas9 mutants.

Specific gRNA and Alt-R™ *S.p.* Cas9 Nuclease V3 were ordered from IDT-DNA and used to assemble in vitro RNP particles; the sequence TCTGCTCCCGCTAAGCCGGC**TGG** in the exon 2 of *Cre16.g665250* (encoding APE1) was selected as the PAM site. *C. reinhardtii* wild-type cells were harvested at early exponential stage ($2\times10^6$ cells·$ml^{-1}$) via centrifugation (1600 g, 5 min, RT), and re-suspended in "MAX Efficiency™ Transformation Reagent for Algae" (Thermo Fisher Scientific), supplemented with 20 % sucrose, to a final concentration of $2\times10^8$ cells·$ml^{-1}$. A heat-shock (40 °C, 30 min, 450 rpm shaking) was then applied to the cells, followed by a 20-min recovery at RT, before 120 µl of cells were added to a pre-cooled 2-mm gap electroporation cuvette together with 15 µl of RNP particles and 1 µg of paromomycin-resistance plasmid for electroporation (600 V, 50 µF, infinite external resistance). Cells were allowed to recover overnight in TAP medium supplemented with 20 % sucrose and 1 mg·$l^{-1}$ Vitamin B12 at 33 °C in the dark before being plated in TAP + 2 % agar + 10 mg·$l^{-1}$ paromomycin and incubated under 20 µmol photons·$m^{-2}$·$s^{-1}$ illumination until colonies could be picked.

DNA isolation and Polymerase Chain Reaction (PCR).

Total genomic DNA and PCR were done using the "Phire Plant Direct PCR Master Mix" kit (Thermo Fisher Scientific) according to the manufacturer's instructions. Briefly, a small amount of cells was solubilized with 20 µl of "Dilution buffer" (included in the PCR kit) via agitation, before centrifugation (4000 g, 10 min, RT) to obtain the DNA in the soluble fraction. The PCR master mix was prepared by mixing 5 µl of 2X reagent with 3 µl of 3 M betaine, 1 µl of each primer, and 1 µl of DNA template (or ddH2O as a negative control), and amplification was done during 35 cycles. PCR products were loaded on a 2 % TAE-agarose gel and the DNA was stained for visualization with ClearSight (Euromedex). The primers used in this manuscript are listed in Supplemental Table 5.

Cloning.

The synthetic DNA sequences of the predicted soluble domain of *C. reinhardtii APE1* (corresponding to the residues Q122 to E276 of *Cre16.g665250*), *A. thaliana APE1* (corresponding to the residues E146 to S286 of *At5g38660*), and *Synechocystis sp. PCC6803 APE1* (corresponding to the residues E54 to P184 of *slr0575*), flanked by complementary *BsaI* restriction sites sequences, were ordered from Twist Bioscience and sub-cloned into the plasmid pLIC03 (LIC:

ligation-independent cloning (*39*) using the GoldenGate cloning technique; the synthetic DNA sequence of *C. reinhardtii PCC1* gene without the predicted chloroplast transit peptide (corresponding to the residues A39 to Q118 of *Cre05.g248600*), flanked by complementary *BsaI* restriction sites sequences, was ordered from Twist Bioscience and sub-cloned into the plasmid pNIC28-Bsa4 (Addgene #26103) using GoldenGate cloning technique. Both pLIC03 (modified Novagen pET-28a+) and pNIC28-Bsa4 vectors contain a 6×His-tag and a TEV protease-cleavage site followed by the suicide gene *SacB* (flanked by *BsaI* restriction sites), which is replaced by our synthetic gene sequences. 20 fmol of plasmid was used with 20 fmol synthetic DNA for three successive rounds of digestion (*BsaI*, 37°C, 10 min) and ligation (T4 DNA ligase, 16°C, 10 min) in the same mix. TOP10 *E. coli* cells (NEB) were then transformed with the ligation products for vector amplification, screening and sequencing. 50 ng of vector were finally used to transform BL21(DE3) cells (Invitrogen) chemically competent *E. coli* for protein expression.

Site-directed mutagenesis.

Site-directed mutagenesis for the different CrAPE1 recombinant versions was based on (*40*). Briefly, pLIC03-APE1 was amplified via PCR using NEB Q5 High-Fidelity 2X Master Mix in 25 µl reaction volumes containing 10 ng DNA template and 0.5 µM of specific primers pairs carrying the new codon (C128S mutation: AGC; H163A: GCG); the PCR protocol was set according to the manufacturer's instructions (amplification was done for 25 cycles with 180 s elongation time each) and annealing temperatures were calculated with the online NEB Tm calculator tool. After amplification, 1 µl of PCR product was incubated at RT for 10 min with NEB 10X KLD Enzyme Mix in a 10 µl reaction volume before being used to transform NEB 5-alpha competent E. coli cells, for plasmid isolation and Sanger sequencing (Eurofins Genomics). Plasmids with the correct sequence were used to transform NEB BL21(DE3) competent E. coli cells for recombinant protein production.

Recombinant protein production.

Bacterial cells were cultured at 37 °C in Terrific Broth Media (APE1) or Luria Broth Media (PCC1) supplemented with 50 mg·l$^{-1}$ of kanamycin (and 25 mg·l$^{-1}$ of chloramphenicol for APE1). Expression was induced once cells reach an OD600 of 0.6 by addition of 0.5mM IPTG, and the cultures were left overnight at 17 °C. Cells were collected by centrifugation and re-suspended in IMAC buffer (50 mM Sodium-Phosphate buffer, 300 mM NaCl, 10 mM Imidazole pH 8.0) supplemented with protease inhibitor cocktail (Roche) and DNase (Sigma-Aldrich). For PCC1-APE1 complex, an equal amount of each pellet was mixed before lysis.

Cells were disrupted twice by French Press at 1000 psi and clarified by centrifugation at 40000 x g for 45 min. The 0.45 µM filtered lysates were loaded on a HiTrap Chelating HP 5ml column (Cytiva), pre-equilibrated with IMAC buffer. Bound protein was eluted with IMAC buffer containing 125 mM imidazole and protein-containing fractions were pooled and dialyzed overnight in 50 mM Sodium-Phosphate buffer, 300 mM NaCl pH 8, with TEV protease in a ratio of 1/20. The resultant was then loaded on another HiTrap Chelating HP 5ml column (Cytiva), pre-equilibrated with IMAC buffer and the untagged protein was collected in the flowthrough. This fraction was then loaded on a HiLoad 16/60 Superdex 75 column, pre-equilibrated in 20 mM MES, 150 mM NaCl, pH 6.5. Fractions containing the target protein were pooled and concentrated using a AMICON centrifugal filter device (Millipore) with a cut-off size of 3.5 kDa. Protein purity was confirmed by SDS–polyacrylamide gel electrophoresis and mass spectrometry. Protein quantification was assayed spectrophotometrically using theoretical extinction coefficient at 280 nm of 8605 $M^{-1}\cdot cm^{-1}$ for APE1 WT and APE1 H43A, 8480 $M^{-1}\cdot cm^{-1}$ for APE1 C8S , 1615 $M^{-1}\cdot cm^{-1}$ for PCC1. Metals were removed post purification by treatment with Chelex or EDTA.

Mass spectroscopy of native proteins.

Proteins were desalted by dialysis using 50mL 20mM ammonium acetate overnight at 4°C. A diluted 200 µL volume was injected at 7µl/min in a Quadripole MS QTOF (Bruker Impact II). The data was recorded using the Otof Control Software then treated using Compass Data Analysis software. After purification, the molecular mass of the proteins corresponded to the predicted mass of the polypeptide for *Cr*APE1 (see Supplemental Table 2), for *Sy*APE1 15114.5Da (theoretical mass: 15115) and for *At*APE1 16024.3Da (theoretical mass : 16025).

Size-Exclusion Chromatography with Multi-Angle Light Scattering.

Multi angle light scattering (MALS) is a technique that determines the precise molecular weight of the species in samples. Knowing that the variation of refractive index related to variation of protein concentration is constant (dn/dC), measuring the refractive index allows one to access the concentration in protein at any point of the elution. 20 µl of protein with a 1.4 $mg.ml^{-1}$ (80 µM) concentration were loaded on an analytical Xbridge Premier Protein SEC 250 Å, 2.5µM column connected to a MALS Dawn 8 spectrometer (Wyatt Instruments). CuCl2 was added to a 1:5 (protein:metal) molar and incubated 5 min with the protein prior to injection. In redox experiments, proteins were first incubated at 25 °C for 2 h with freshly prepared 5 mM $H_2O_2$ or for 10 min with 1 mM tris(2-carboxyethyl)phosphine (TCEP). The column was pre-equilibrated at 0.6 ml/min in 20 mM MES, 150 mM NaCl, pH 6.5 buffer filtrated at 0.1 µm. An in-lane refractive index detector (Optilab, Wyatt Instruments) was used to follow the differential refractive index relative to the solvent. After baseline correction, all samples presented isolated peaks allowing the determination of absolute molecular masses using ASTRA6 software (Wyatt Instruments) and a theoretical dn/dc value of 0.185 ml/g.

Thermal unfolding experiments by nano differential scanning fluorimetry.

We tested the binding of various divalent cations by nano differential scanning fluorimetry, that measures the quenching (at 330 nm) and red-shifting (at 350 nm) of the protein fluorescence arising from binding-induced changes in the environment (from solvent to protein moiety) of residues, mainly tryptophan. For thermal unfolding experiments, APE1 proteins were diluted to a final concentration of 40 µM and incubated with 1 molar excess of each divalent metal or with a range of $CuCl_2$ from 1 to 200 µM. For each condition, 10 µl of sample per capillary were required for one thermal unfolding profile. The samples were loaded into Prometheus capillaries and experiments were carried out using the Prometheus NT.48 (NanoTemper Technologies, GmbH). The temperature ramp was set to an increase of 2 °C/min in a range from 25 °C to 90 °C. Protein unfolding was measured by detecting the temperature-dependent change in tryptophan fluorescence at emission wavelengths of 330 and 350 nm. For calculation of Melting Temperature (Tm), the first derivative data of the F350/F330 fluorescence ratio unfolding curves were used.

Protein interaction by MicroScale Thermophoresis.

PCC1 was diluted to 17µM and labelled using Monolith Protein Labeling Kit RED-NHS 2nd generation (NanoTemper Technologies GmbH). For APE1, a serial 1:1 dilution was performed by transferring 10 µL of 5 µM APE1 solution to an equal volume of working buffer (20 mM MES, 150 mM NaCl, pH 6.5) mixing, and repeating this step 16 times. This way, the ligand concentration is reduced by 50% in each dilution step. 10µl of labelled PCC1 at 34nM was mixed with 10µL of each APE1 serial dilution from 2.5 to 7,7.10-5 µM. For the quantification of copper binding to APE1, a serial dilution of $CuCl_2$ from 1.6 $10^{-5}$ M to 4.88 $10^{-10}$ M was prepared in 20 mM MES, 150 mM NaCl, pH 6.5. Labelled APE1 was added to a final concentration of 24 nM. Labelling of APE1 has to occur just before MST experiments because it strongly impairs APE1 stability. MST experiments were performed on a NanoTemper ® Monolith NT.115 with red filter (NanoTemper Technologies GmbH). Samples were loaded into standard treated capillaries. Measurements were performed at 25 °C using 100 % excitation on medium MST power position. Data analysis was performed using the NanoTemper® analysis software, where $K_D$ constants was calculated using the saturation binding curve at equilibrium.

Superoxide detoxification assays.

Superoxide detoxification activity was evaluated by a specific assay kit (CS0009, Sigma-Aldrich), according to the manufacturer's instructions or diluted 4 times. Superoxide detoxification activity was proportional to the decrease in the color signal of WST-1 (Water Soluble Tetrazolium dye) due to $O_2^{\bullet-}$ production by xanthine/xanthine oxidase, determined by reading the absorbance at 450 nm wavelength after a given reaction time at 25 °C (Infinite M200 spectrophotometer, Tecan Ltd) after subtracting the background (buffer control). Molar extinction coefficient of WST-1 at 450 nm is $\varepsilon_{450nm} = 35.2 mM^{-1}.cm^{-1}$ (*41*). In the 96 well plate with 200 µl assay volume the optical path is 0.5 cm. In the experimental conditions of Fig. 4C, absorbance increased by 0.1 (O.D.) every 20 min, corresponding to 5.7 µM formazan, each formazan molecule quenching two $O_2^{\bullet-}$ molecules. The rate of generation of superoxide radical anions was therefore 0.57 $\mu M.min^{-1}$. In Fig. 4D the rate of generation was calculated as 1.14 $\mu M.min^{-1}$

EPR spectroscopy.

X-band continuous wave EPR spectra were recorded with a Bruker Elexsys 500 X-band spectrometer equipped with a standard ER 4102 (Bruker) X-band resonator, a Bruker teslameter, an Oxford Instruments cryostat (ESR 900) and an Oxford ITC504 temperature controller. Spectra were collected at 30 K; modulation amplitude, 10G; microwave power, 0.02 mW; microwave frequency, 9.5 GHz; modulation frequency, 100 kHz. 50µM protein samples were suspended in 100mM HEPES pH 6.5 with equimolar concentrations of $CuCl_2$. Spectra were fitted using EasySpin, version 6 (*42*).

Metal determination and quantification**.**

Metal content of the proteins (Cu, Zn) was quantified by MP-AES (microwave-plasma-atomic-emission-spectroscopy). Samples were treated with 50% nitric acid and heated to 96C° for 2h then diluted to 10% nitric acid prior measurement. After 10-fold dilution in trace-metal-free water, the metal content of the samples was determined by atomic emission spectroscopy using an MP AES 1200 spectrometer (Agilent, USA).

Statistical analysis.

The statistical relevance was calculated using GraphPad Prism 10 by one-way Analysis of variance (ANOVA), a comparison between the means of two or more groups by analyzing variance, using Tukey's post-hoc multiple comparisons, and shown as pairwise comparisons (p≤0.001*** or p≤0.0001 ****) or compact letter display (CLD).

Experimental set up for measuring plastocyanin accumulation *in vivo*

The protocol was adapted based on (*43*). Cells were spun down and resuspended in Ficoll and dark adapted. Cells were then poisoned with DCMU and DBMIB to restrict electron flow to only two secondary electrons donors to photo-oxidized $P_{700}^{+}$ in WT: cyt *f* and PC. Flash-induced absorbance changes were measured using the Joliot Type Spectrophotometer (JTS-10) using two detection wavelengths (705 nm for $P_{700}^{+}$, and 740 nm to subtract $PC^{+}$ contribution). Saturating single turnovers of PSI were induced by a < 10 ns laser flash three biological replicates were measured.

Analysis of genes co-expressed with *APE1*

Co-expression analyses were performed to identify genes coordinately expressed with the three APE1 orthologs chosen in this work. For Chlamydomonas reinhardtii *APE1* (Cre16.g665250), the PhytoMine expression compendium tool of JGI Phytozome v14 (https://phytozome-next.jgi.doe.gov/) was queried; genes exhibiting a Pearson correlation coefficient (PCC) ≥ 0.80 with *CrAPE1* across 518 public RNA-seq datasets were retained. For Synechocystis sp. PCC 6803 *APE1* (SLR0575), the AlcoDBcyano portal was used to retrieve the top-ranking co-expression partners; only genes with a mutual-rank (MR) ≤ 50 were kept. For Arabidopsis thaliana *APE1* (AT5G38660), co-expression data were obtained from ATTED-II v12.0, and genes with Logit Score > 3.0 (≈ PCC ≥ 0.70) were selected. Genes were manually assigned to functional categories according to their primary annotation; when no annotation was available, assignments were based on additional information obtained from the KEGG database or from targeted literature searches.

**Supplementary Figures.**

**SF1.**

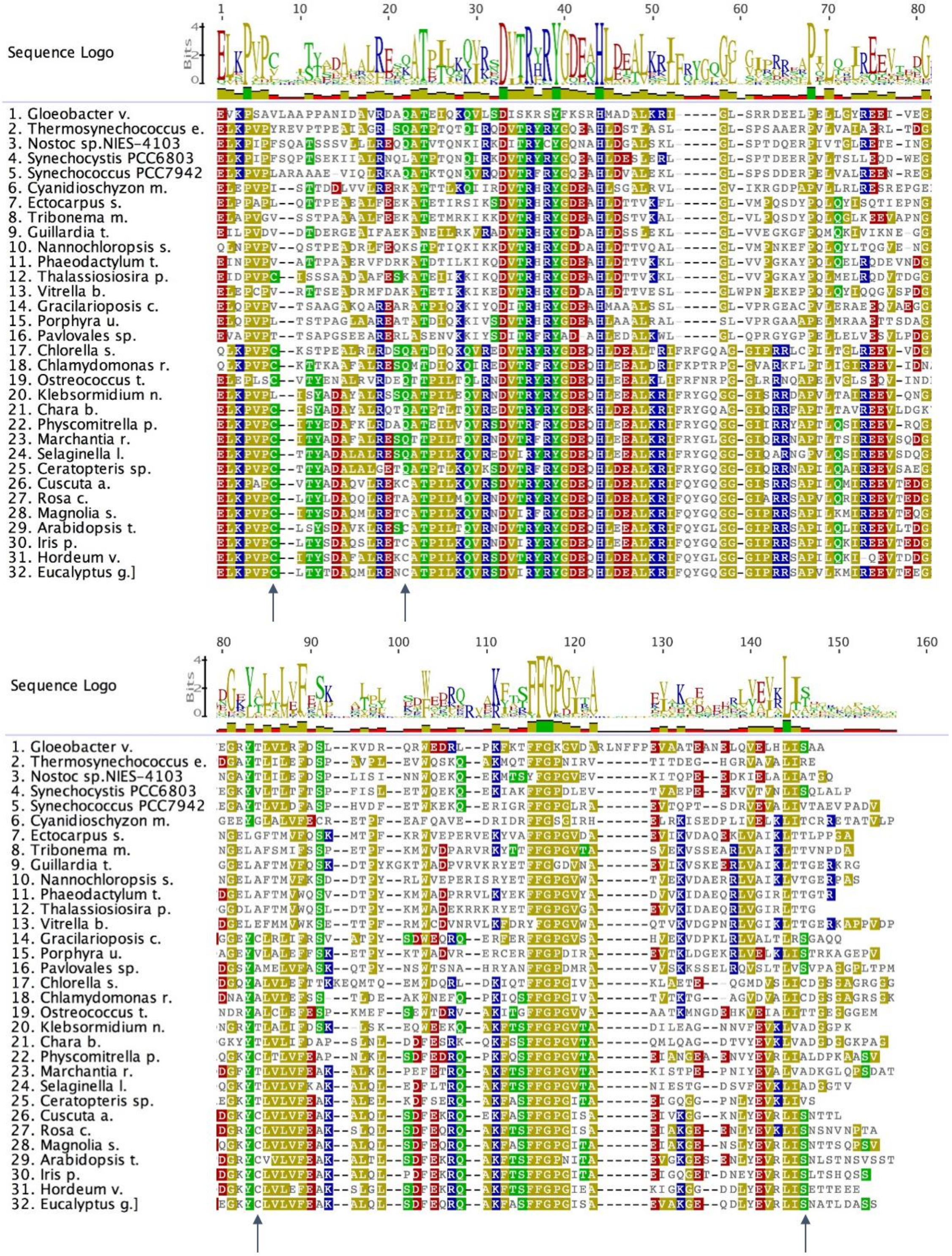


**Supplemental Figure 1**. **Alignment of soluble region of APE1 expressed a recombinant protein from major representatives of oxygenic phototrophs** MASSF Alignment of soluble region of APE1 (part expressed as a recombinant protein) from major representatives of oxygenic phototrophs: cyanobacteria

(1-5), red algae (6,14,15), haptophytes, heterokonts (7, 8, 10,11,12), chromelladiophytes (13), cryptophytes (9), haptophytes (16) chlorophytes (17,18,19), charophytes (20,21) bryophytes (22,23), lycophytes (24,25) and different land plants (26-32). Variable cysteines are shown with an arrow.

**SF2.**

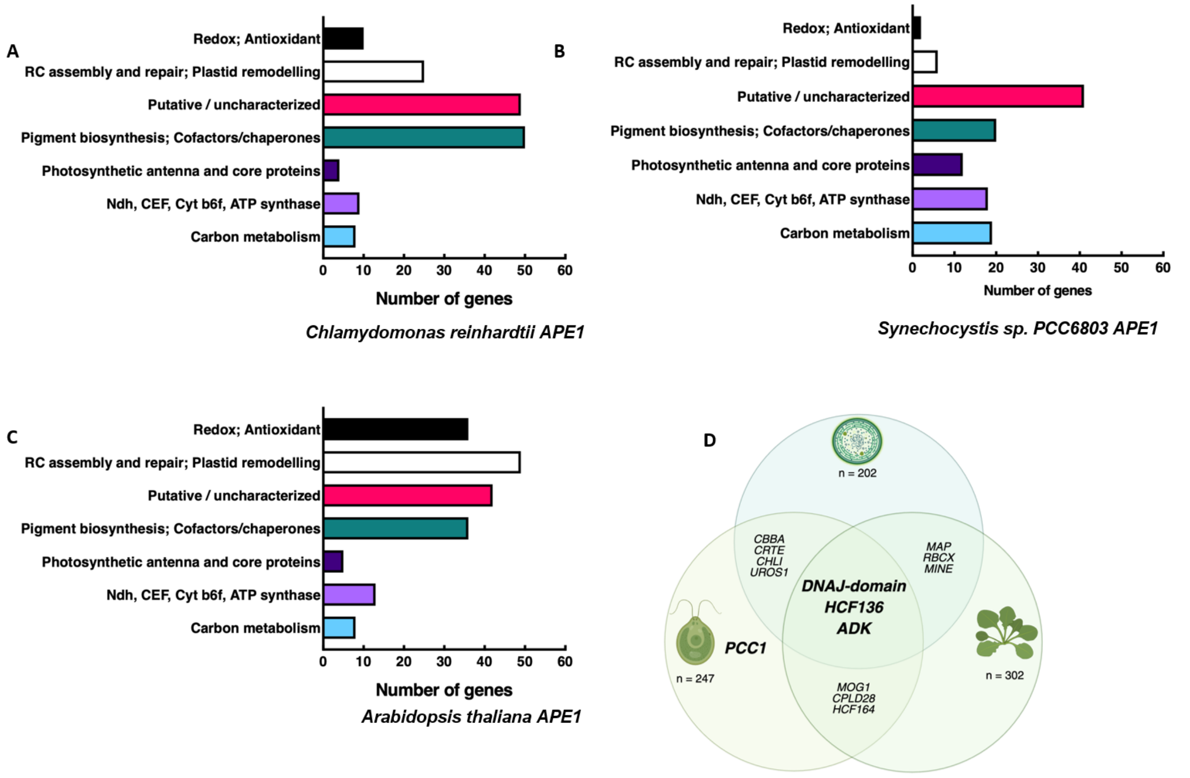


**Supplemental Figure 2. Data mining of *APE1* co-expressed genes supports a similar function across oxygenic phototrophic species.** *APE1* from A. *Chlamydomonas reinhardtii*, B. *Synechocystis sp PCC6803*, and C. *Arabidopsis thaliana* share similar co-expressed genes classed into categories related to biogenesis of the photosynthetic apparatus and photoprotection. D. A small subset of orthologous genes show shared co-expression between model species: Chlamydomonas and Synechocystis tables both included *CBBA* (encoding Fructose-1,6-bisphosphate aldolase by *CRE02.G093450* or *SLL0018*), *CRTE* (encoding Geranylgeranyl pyrophosphate synthase by *CRE01.G050950* and *SLR0739*), *CHLI* (encoding Magnesium protoporphyrin IX chelatase subunit I by *CRE06.G306300* or *SLR1030*), and *UROS1* (encoding Uroporphyrinogen-III synthase by *CRE09.G409100* or *SLR1887*); Chlamydomonas and Arabidopsis tables both include *MOG1* (encoding a FKBP-type peptidyl-prolyl cis-trans isomerase by *CRE10.G454250* or *AT1G77090*), *CPLD28* (encoding LOW PSII ACCUMULATION 3 by *CRE03.G184550* or *AT1G73060*), and the thioredoxin *HCF164* (*CRE17.G702150* or *AT4G37200*); Arabidopsis and Synechocystis tables both include *MAP* (encoding Methionine aminopeptidase by *AT4G37040* or *SLL0555*), *RBCX* (encoding a putative Rubisco chaperonin by *AT1G69390* or *SLR0011*), and *MINE* (encoding a cell division factor by *AT1G69390* or *SSL0546*); all three tables include genes encoding DNAJ-domain proteins (for example *AT5G03880*, *CRE07.G316050*, and *SLL1933*), *HCF136* (*CRE06.G273700*, *SLR2034*, or *AT5G23120*), and ADK (encoding Adenylate kinase by *CRE17.G739550*, *SLL1815*, or *AT5G35170*). Panel D. was created using BioRender.com. All co-expressed genes are shown in supplemental data set 1.

**SF3.**

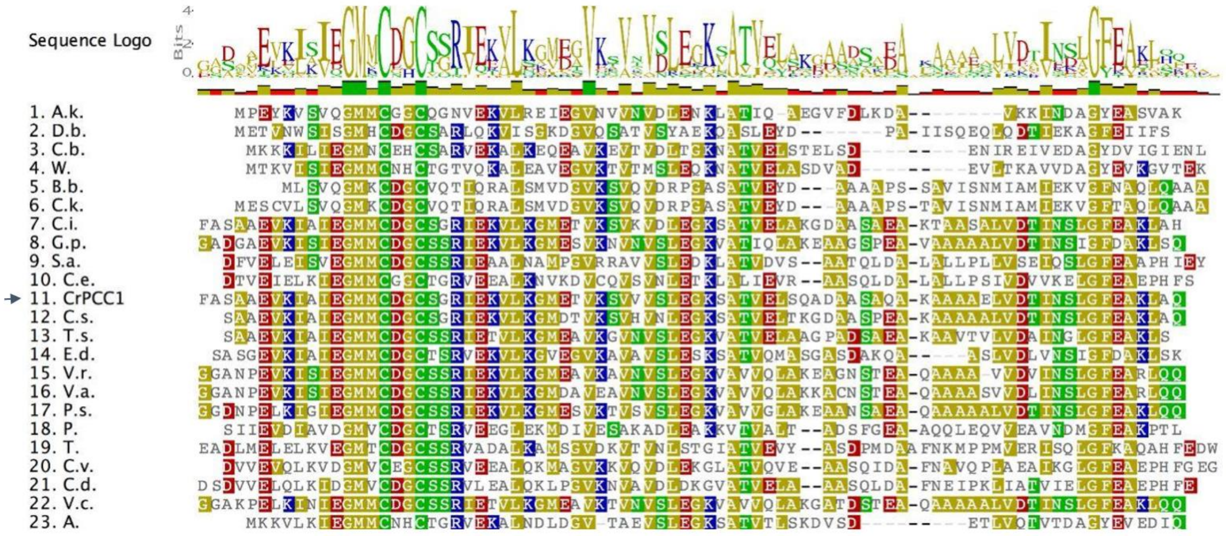


**Supplemental Figure 3. Alignments of CrPPC1 orthologues** (marked with an arrow). Orthologues of CrPCC1 were identified in the *Chlamydomonales*. The Chlorophyceae, Trebouxiophyceae and different aerobic and anaerobic bacteria isolated from different environments and one amoeba also have a protein resembling PCC1. *A.k., Acrasis kona (soil dwelling amoeba); D.b., Deltaproteobacteria bacterium (myxococcota, fruiting, gliding, gram -ve); C.b., Clostridia bacterium (gram +ve bacteria); W., Waltera sp. (intestinal bacteria); B.b., Bacteroidetes bacterium (gram -ve gut bacteria, aerobic or anaerobic); C.k. , Candidatus Kapaibacterium sp. (green sulfur, anaerobic phototroph); C.i., Chlamydomonas incerta (Chlamydomonales. Chlorophyceae); G.p. , Gonium pectorale (Chlamydomonales, Chlorophyceae, colonial); S.a., Sanguina aurantia (Chlamydomonales, Chlorophyceae, colonial, red snow); C.e., Chlamydomonas eustigma (Chlamydomonales, Chlorophyceae); CrPCC1, Chlamydomonas reinhardtii (Chlamydomonales, Chlorophyceae); C.s., Chlamydomonas schloesseri (Chlamydomonales, Chlorophyceae); T.s., Tetrabaena socialis (Chlamydomonales, Chlorophyceae); E.d., Edaphochlamys debaryana  (Chlamydomonales, Chlorophyceae); V.r., Volvox reticuliferus  (Chlamydomonales. Chlorophyceae); V.a., Volvox africanus (Chlamydomonales, Chlorophyceae); P.s., Pleodorina starrii  (Chlamydomonales. Chlorophyceae, colonial); P., Picochlorum sp. BPE23 (Trebouxiophyceae); T., Trebouxia sp. C0004 (Trebouxiophyceae); C.v., Chlorella variabilis (Trebouxiophyceae); C.d. ; Chlorella desiccata (Trebouxiophyceae); V.c., Volvox carteri f. Nagariensis (Chlamydomonales, Chlorophyceae); A.,  Anaerotignum sp. (strict  anaerobe bacteria)*

**SF4.**

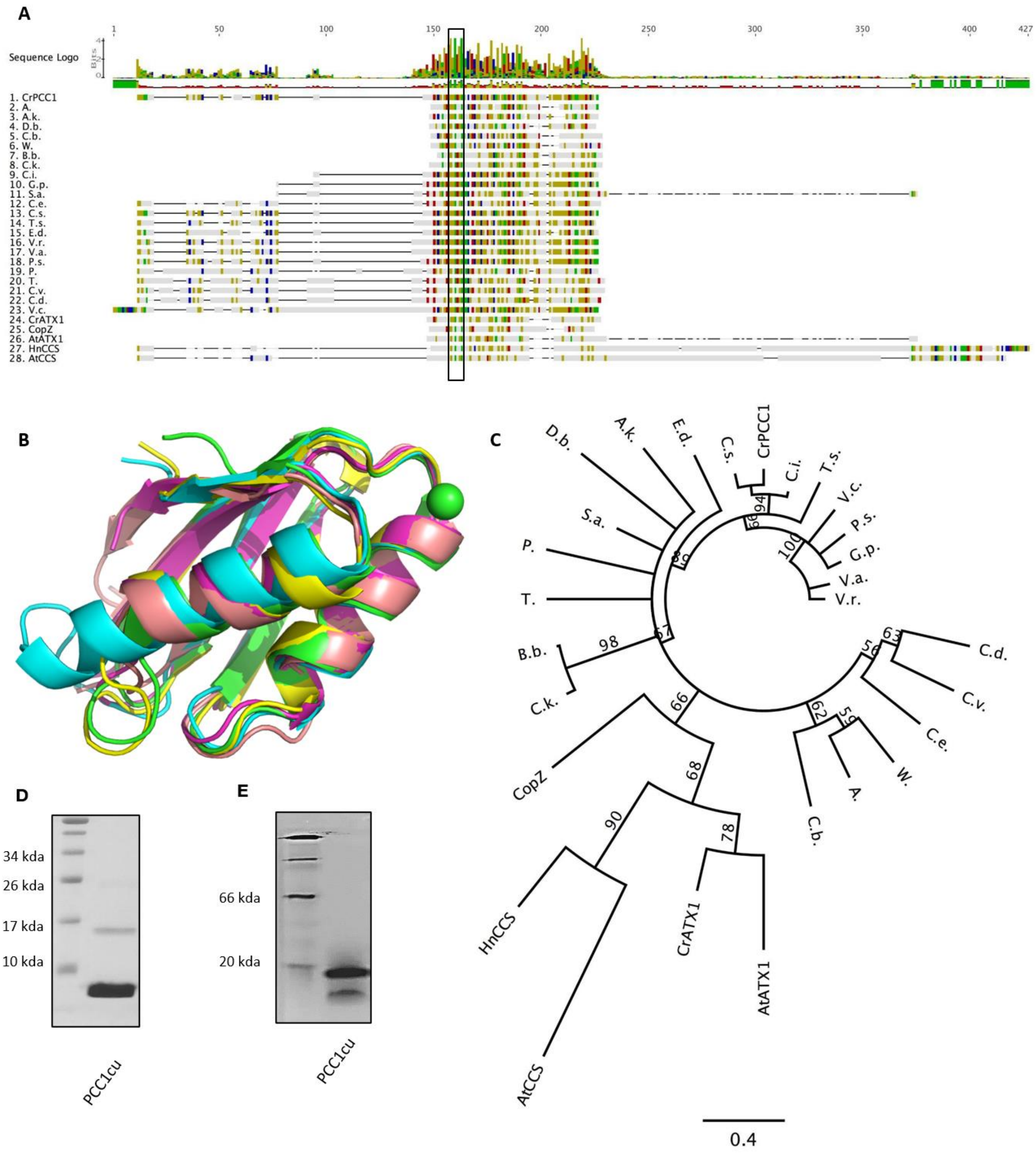


**Supplemental Figure 4. PCC1 can be considered ATX1-like, its structure shows the classic ferredoxin-like folds and HMBD, but is not a true orthologue of ATX1. PCC1 orthologues are restricted to a handful of green algae, but appear to be widespread in bacteria A.** Alignments showing coloured zones of high conservation between orthologues of PCC1, followed by orthologues of ATX1 including CCS1 orthologues. The Heavy Metal Binding Domain (HMDB) MxCxxC is boxed. B. The Alphafold3 predicted structure of Cr Plastid Cu Chaperone 1, PCC1 [A0A2K3DRZ6] (cyan), shows high structural homology to: BsCopZ loaded with Cu2+ [O32221] (X-ray structure, green), cytosolic CrATX1 [9FYV4] (predicted structure, magenta), domain I of CrCTP3 [A0A2K3DF91] (predicted structure, yellow), and domain 1 of Hn CuZnSOD chaperone CCS1[O14618] (X-ray structure, tint); copper ion is represented as a green sphere. C. Evolutionary relatedness of ATX domain containing PCC1 orthologues and structural orthologues. D. The purified recombinant PCC1 is a 8kDa protein with traces at 16kDa dimer when separated via semi-denaturant SDS-PAGE. E. PCC1 forms a 16 kDa dimer when separated by native-PAGE conditions. *Abbreviations: A., Anaerotignum sp. (strict anaerobe bacteria); A.k., Acrasis kona (soil dwelling amoeba); D.b., Deltaproteobacteria bacterium (myxococcota, fruiting gliding, gram -ve); C.b., Clostridia bacterium (gram +ve bacteria); W., Waltera sp. (intestinal bacteria); B.b., Bacteroidetes bacterium (gram -ve gut bacteria, aerobic or anaerobic); C.k. , Candidatus Kapaibacterium sp. (green sulfur, anaerobic phototroph);* ***C.i., Chlamydomonas incerta (Chlamydomonales. Chlorophyceae);*** *G.p. , Gonium pectorale (Chlamydomonales, Chlorophyceae, colonial); S.a., Sanguina aurantia (Chlamydomonales, Chlorophyceae, colonial, red snow); C.e., Chlamydomonas eustigma (Chlamydomonales, Chlorophyceae);* ***CrPCC1, Chlamydomonas reinhardtii (Chlamydomonales, Chlorophyceae); C.s., Chlamydomonas schloesseri (Chlamydomonales, Chlorophyceae);*** *T.s., Tetrabaena socialis (Chlamydomonales, Chlorophyceae); E.d., Edaphochlamys debaryana (Chlamydomonales, Chlorophyceae); V.r., Volvox reticuliferus (Chlamydomonales. Chlorophyceae); V.a., Volvox africanus (Chlamydomonales, Chlorophyceae); P.s., Pleodorina starrii (Chlamydomonales. Chlorophyceae, colonial); P., Picochlorum sp. BPE23 (Trebouxiophyceae); T., Trebouxia sp. C0004 (Trebouxiophyceae); C.v., Chlorella variabilis (Trebouxiophyceae); C.d. ; Chlorella desiccata(Trebouxiophyceae); V.c., Volvox carteri f. Nagariensis (Chlamydomonales, Chlorophyceae); CrATX1, Chlamydomonas reinhardtii ATOX1 orthologue; CopZ, bacillus subtilis; HnCCS, Human copper chaperone for CuZn SOD; AtATX1, Arabidopsis thaliana ATOX1; AtCCS, Arabidopsis thaliana copper chaperone for CuZn SOD.*

**SF5.**

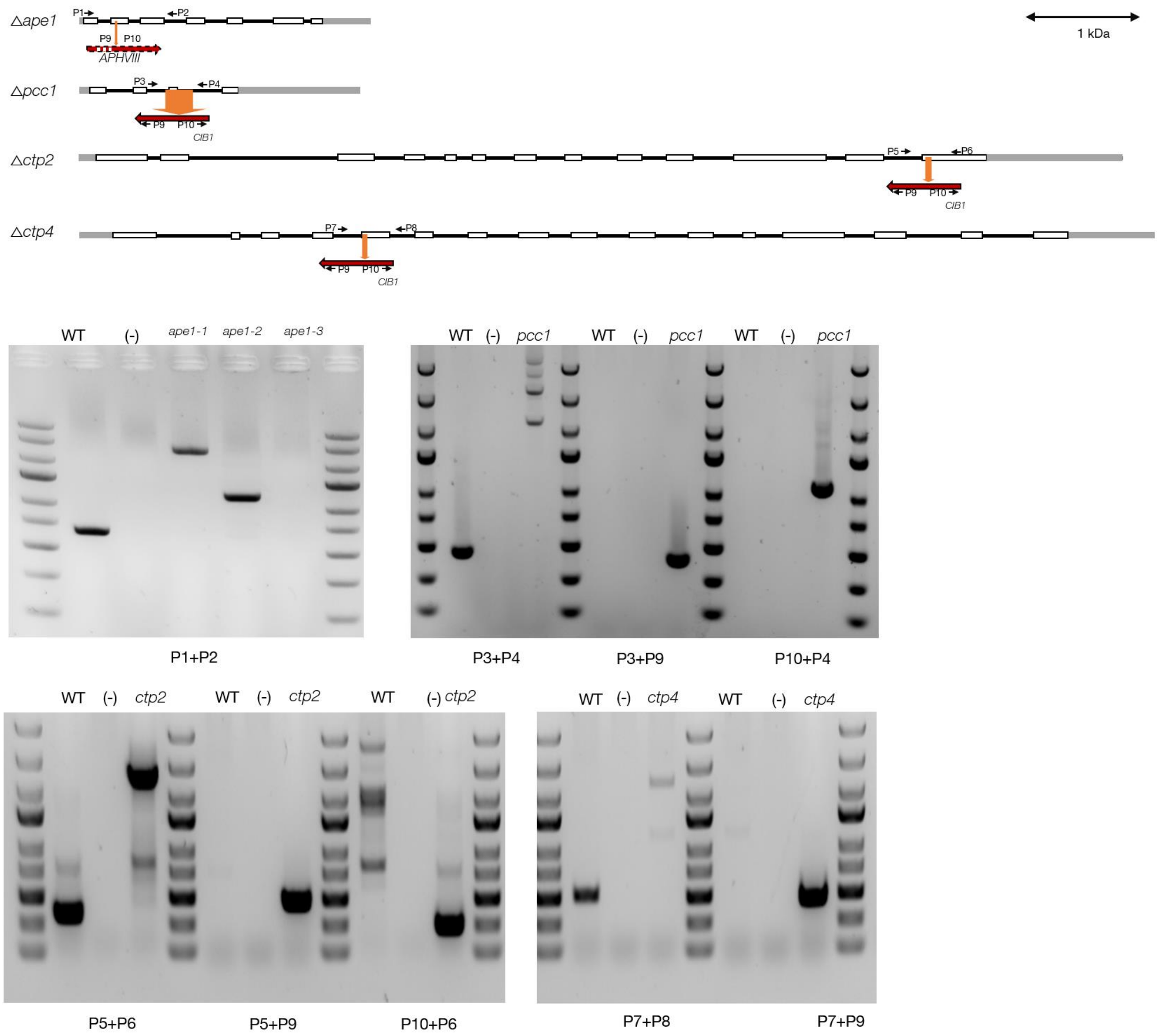


**Supplemental figure 5. Confirmation of the predicted *C. reinhardtii* insertions sites in the different lines used in this study via PCR** (details regarding the primers used can be found in Supp. table 3). The WT strain is T222 (for *ape1*) or CC-5325 (MJ CLiP mutant collection background). *Acclimation of Photosynthesis to the Environment 1*, *ape1*, mutants contain an insertion with variable size in exon 2 (upon targeting via CRISPR/Cas9); Plastid Cu Chaperone 1, *pcc1*, mutant contains an insertion from intron 2 to intron 3, with deletion of exon3 *(LMJ.RY0402.059810, Cre05.g248600); Cu transporter 2*, *ctp2*, mutant contains an insertion in exon 13 *(LMJ.RY0402.149111, Cre10.g424775); Cu transporter 4*, *ctp4*, contains an insertion in exon 5 *(LMJ.RY0402.047261, Cre10.g422201*). Promoters and terminators are represented as grey boxes, exons as white boxes, and introns as a black line.

**SF6.**

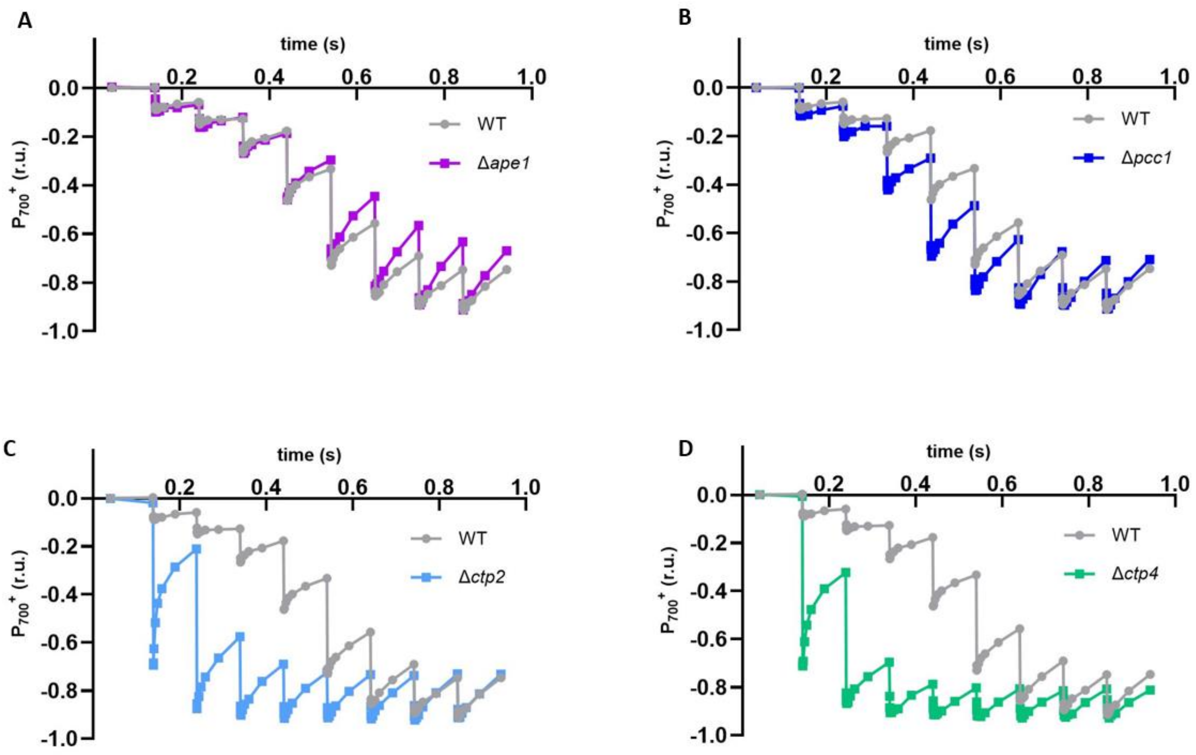


**Supplemental Figure 6. Quantification of secondary electron donors to $P_{700}^{+}$ in intact cells of WT, *pcc1*, *ape1*, *ctp2* and *ctp4* mutants treated with HA, DCMU and DBMIB.** $P_{700}$ absorbance changes at 705 nm was measured following a train of single turnover saturating flashes. In the WT (grey), a positive charge accumulates on $P_{700}$ after 4 to 5 flashes, corresponding to 3 to 4 electrons transferred from Rieske protein, cyt f, and 1 to 2 plastocyanins. While no significant difference was found for *ape1* (purple) **A.** and *pcc1* mutants (royal blue) B. *ctp2* (pale blue) **C**.   and *ctp4* (green) D.  showed that $P_{700}$ was almost fully oxidized on the first flash, showing absence of plastocyanin in these 2 mutants.

**SF7.**

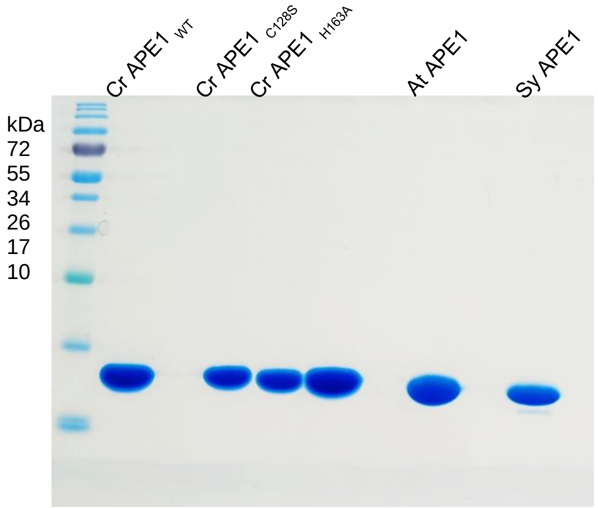


**Supplemental Figure 7. Production of APE1 recombinant proteins.** Denaturing gel SDS-page 15% acrylamide. 10µl of sample post gel filtration at 50 µM are loaded in each well.

**SF8.**

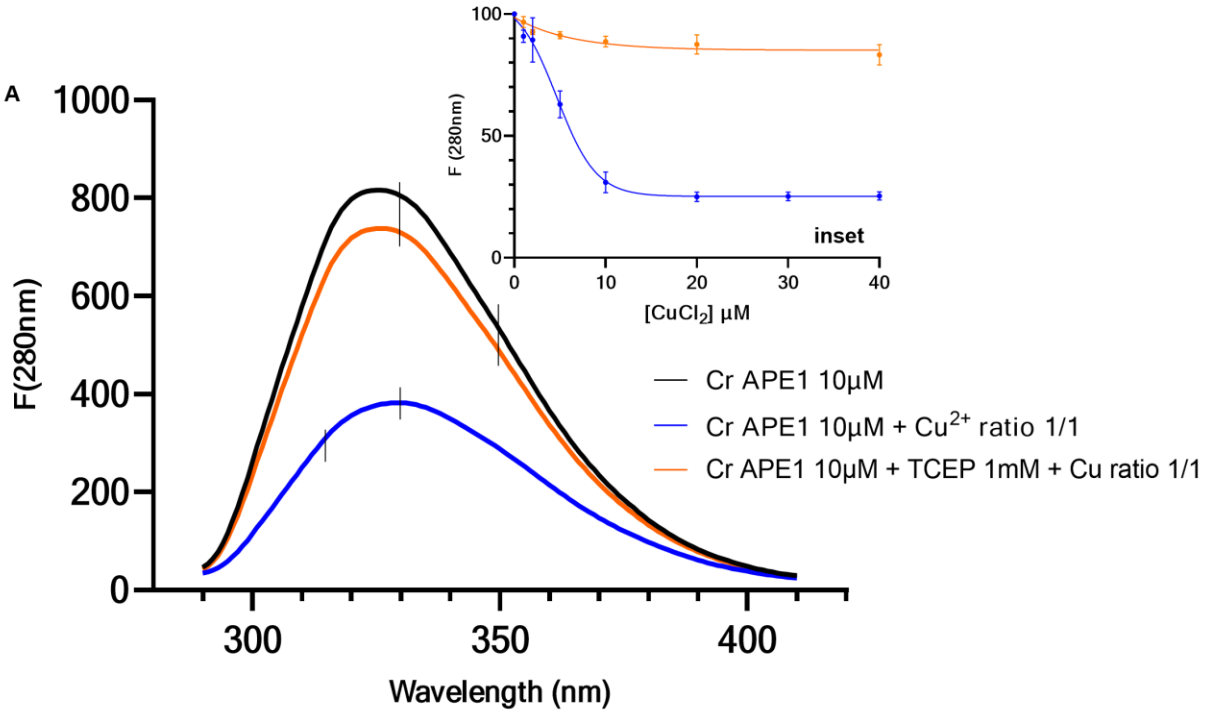


**Supplemental Figure 8. APE1 fluorescence spectra shows that $CuCl_2$ quenches protein (Trp) fluorescence (at 330 nm) and red-shifts (to 350 nm) but this is strongly inhibited in the presence of TCEP. A.** Fluorescence emission spectra (excitation 280 nm) for 10µM APE1 WT (black), 10µM APE1 WT with a molar ratio of Cu(II) (blue) and 10µM APE1 WT incubated 10 minutes with 1mM TCEP then with a molar ratio of Cu(II) (orange). B. Titrations of APE1 WT (10µM) with (orange) or without (blue) 1mM TCEP by increasing concentrations of Cu(II).

**Fig. S9.**

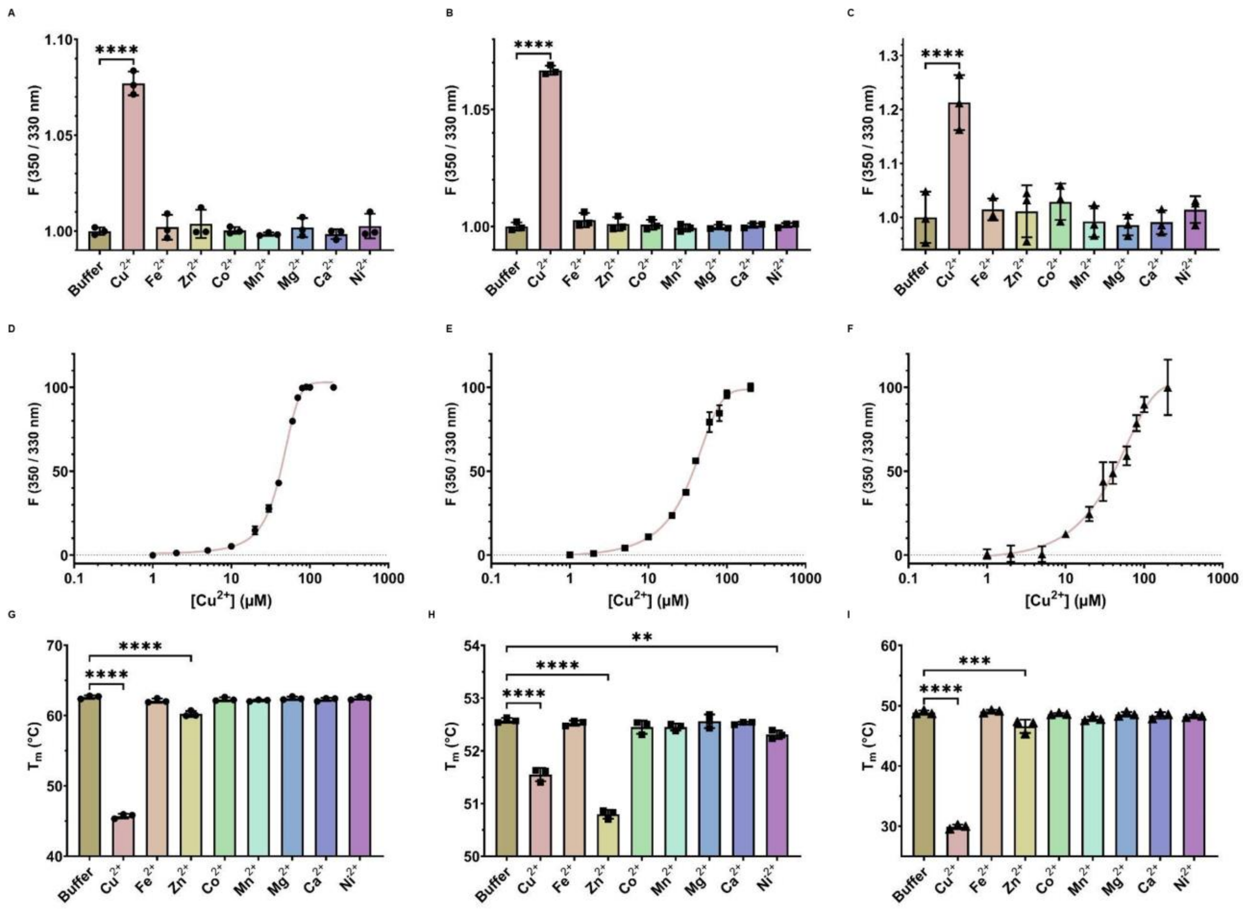


**Supplemental Figure 9. APE1 has a specificity for Copper Binding across different oxygenic phototrophs.** APE1 Recombinant proteins of Chlamydomonas (circles, **A**, D and G) Synechocystis PCC 6803 (squares, B, E and H), and Arabidopsis (triangles, C, F and I) in interaction with different divalent cations. Only $Cu^{2+}$ significantly changes local protein conformation in the environment of Trp measured as an increase in the 350/330 nm fluorescence ratio (A, B and C). Titrations of recombinant APE1 (40µM) with increasing concentrations of $Cu^{2+}$ shows a dose dependant increase in fluorescence (D, E and F). The melting temperature ($T_m$) is calculated from the melt curve; changes in $T_m$ are correlated to changes in protein stability or ligand binding. Both $Cu^{2+}$ and $Zn^{2+}$ significantly change the melting temperature of all recombinant APE1 versions (G, H and I). Titrations were fitted to a concentration-dependent sigmoid model. The statistical relevance is shown as pairwise comparisons ($p \leq 0.0001$ ****, $p \leq 0.001$ *** or $p \leq 0.01$ **).

**SF10.**

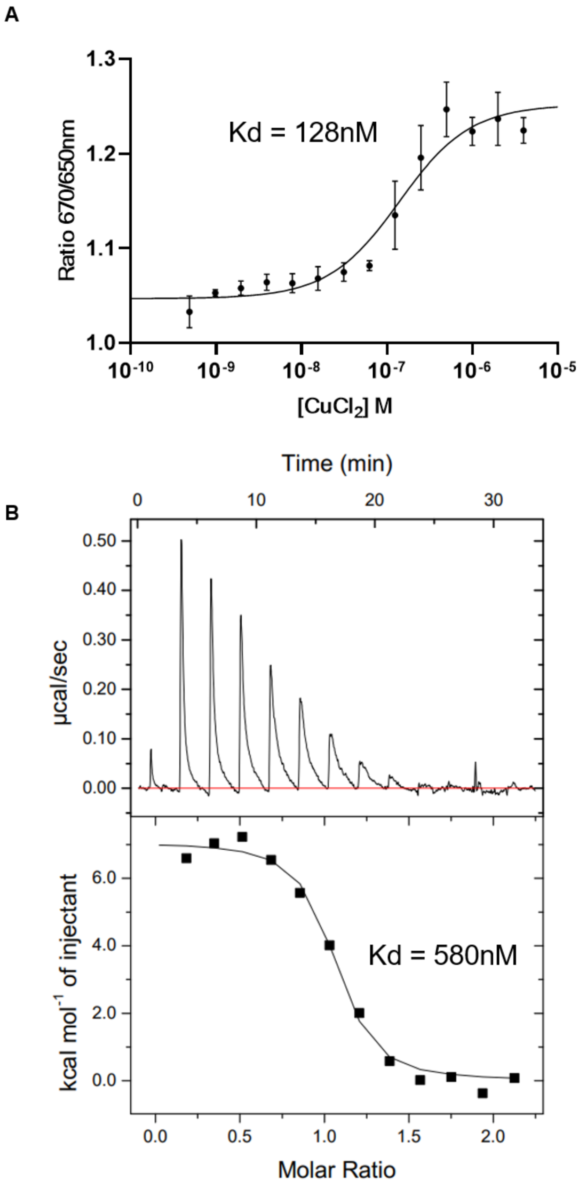


**Supplemental Figure 10. CrAPE1 has high affinity for Cu(II) showing a specific interaction with a dissociation constant ($K_D$) in the nmolar range using 2 different techniques. A.** Microscale Thermophoresis (MTP- Monolith) titrations of fluorescently labelled *Cr*APE1 (24nM) against increasing concentrations of Cu(II) show conformational changes linked to Cu binding, giving a $K_D$ = 128nM (confidence interval 79-206nM). B. Isothermal Titration Calorimetry (ITC) titrations of Cu(II) against 50µM CrAPE1 shows exothermic binding of Cu to CrAPE1 with calculated $K_D$ = 580nM (confidence interval 450 à 840nM).

**SF11.**

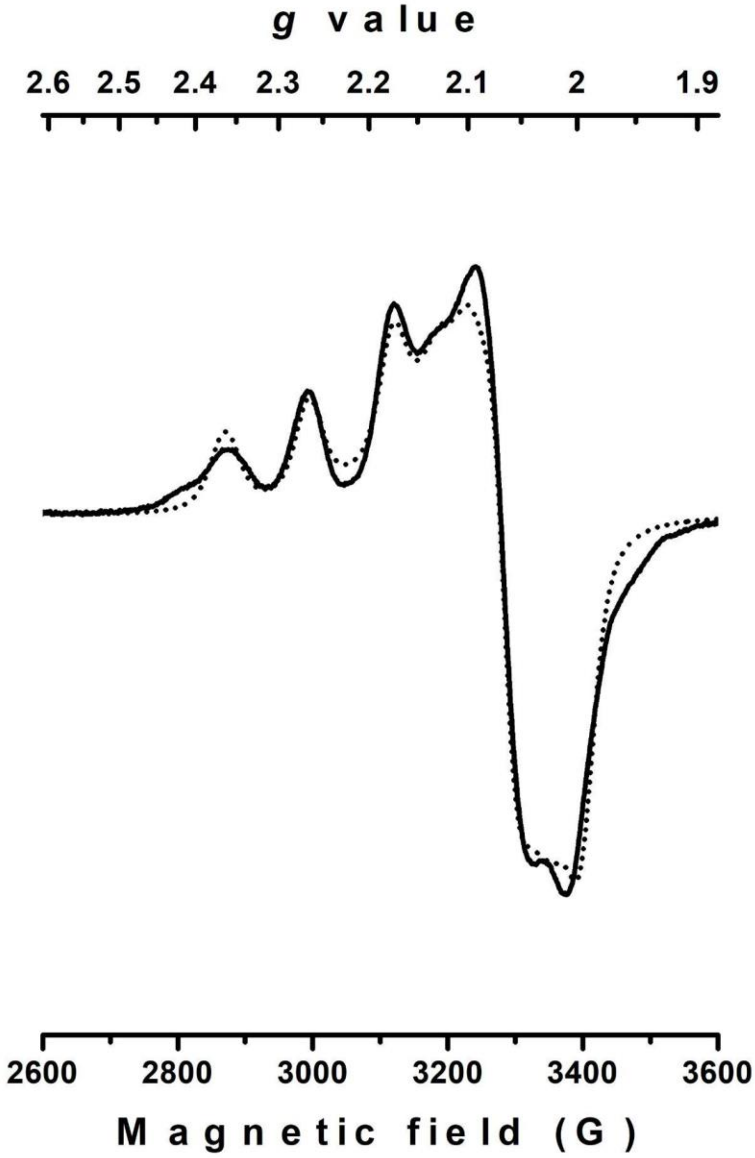


**Supplemental Figure 11. The Fit of X-field EPR spectra of $Cu^{2+}$APE1$_{WT}$** (solid line) against the fit of this spectrum with one component (dotted line) gave *g*-values of $g_z$ = 2.219, $g_y$ = 2.103, and $g_x$ = 2.020, with a hyperfine coupling constant $A_{zz}$ of 378 MHz (134 G). The fit of APE1's spectrum indicated that there were at least two different components. The main component (fitted in Supplemental Figure 11) had principal *g*-values of $g_z$ = 2.219, $g_y$ = 2.103, and $g_x$ = 2.020, with a hyperfine coupling constant $A_{zz}$ of 378 MHz (134 G) indicative of a rhombic environment around the Cu atom. This is however an unusual combination of $g_z$ and $A_{zz}$ constants reflecting an atypical coordination sphere for Cu(II).

SF12.

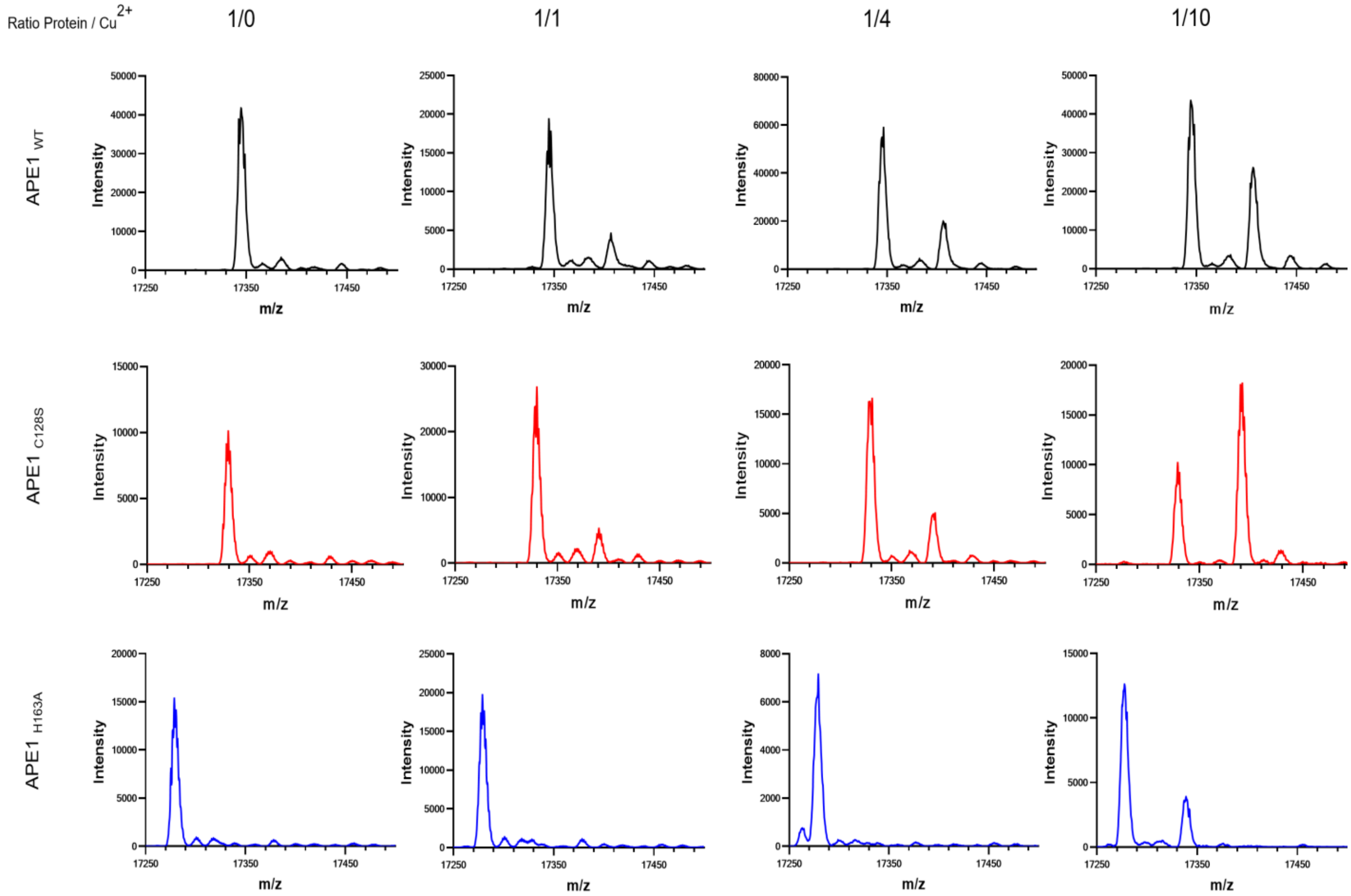


**Supplemental Figure 12.** Native Mass Spectrometry (native-MS) to assess Cu-binding. Mass to charge ration (m/Z) gives molecular weight and sperates apo-APE1 from Cu loaded APE1. Cu(II) binds to $APE1_{WT}$ and $APE1_{C128S}$ at low concentrations (1/1), but only at very high concentrations (1/10) to $APE1_{H163A}$.

**SF13.**

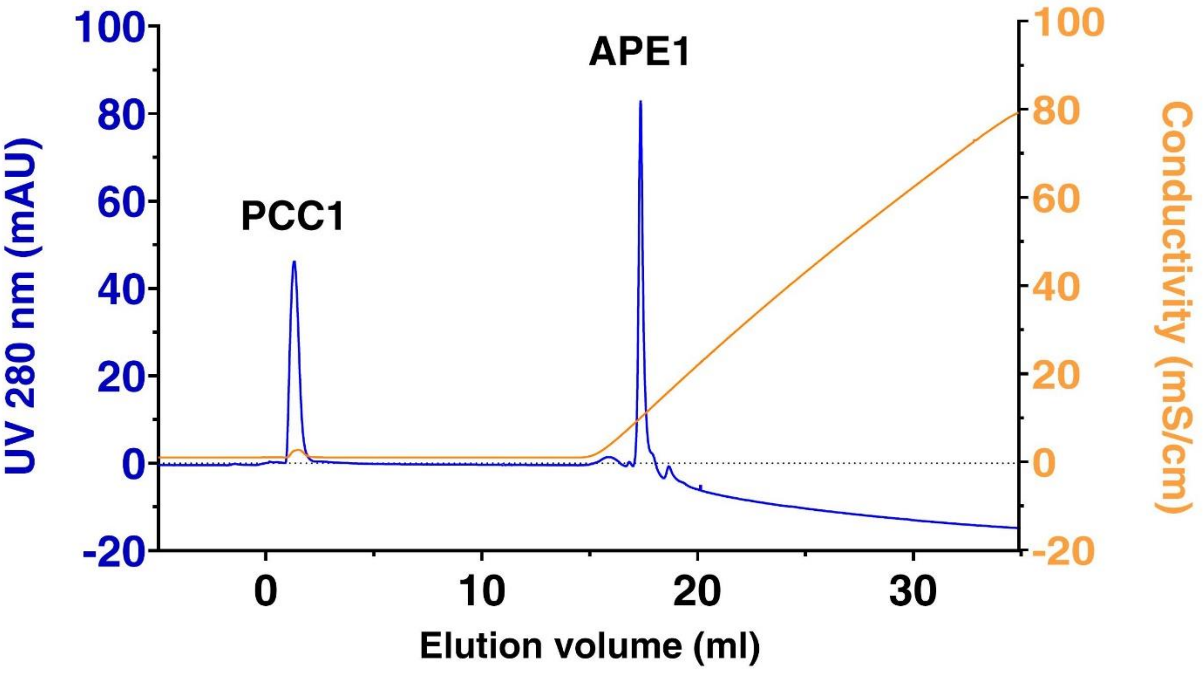


**Supplemental Figure 13**. Separation of APE1-PCC1 complex, 150 mM pre-dialysis, via anion-exchange chromatography (Mono-S column). Fractions were collected, containing variously diluted PCC1 and APE1 quantities around the times of peak identification. Samples were used for AES-MS.

### Supplementary Tables

#### Table S1.

| Annotation | *Synechocystis sp. PCC6803* |
|---|---|
| *PsbB* (CP47) | Slr0906 |
| *PsbC* (CP43) | Sll0851 |
| *PsbE* (α-b559) | Ssr3451 |
| *PsbF* (β-b559) | Smr0006 |
| *PsbH* | Ssl2598 |
| *PsbO* (33kDa) | Sll0427 |
| *PsbP-like* (24kDa) | Sll1418 |
| *APE1* | Slr0575 |
| *TLP15* | Sll1071 |

**Supplementary Table 1 . Genes found in all photosynthetic organisms that have retained PSII.** List of cyanobacterial genes conserved across all oxygenic phototrophs based on a cross comparison of supplementary files of Mulkidjanian et al., 2006 (*22*) and Beck et al., 2018 (*23*) with reference to Zehr et al., 2008 (*24*). The list comprises genes and their proteins, in parentheses, that are always but exclusively found together with the *PsbA* (D1) and *PsbD* (D2) core subunits of PSII, of which they form the reaction centre.

**Table S2.**

| Protein | Ratio Protein / Cu(II) | Theoretical Mass (Da) | Identified Mass (Da) | Attribution |
|---|---|---|---|---|
| ***Cr*APE1$_{WT}$** | - | 17346 | 17345 | apo *Cr*APE1 |
| ***Cr*APE1$_{WT}$** | 1:1 | | 17345<br>17407 | apo *Cr*APE1<br>*Cr*APE1-Cu |
| ***Cr*APE1$_{WT}$** | 1:4 | | 17345<br>17407 | apo *Cr*APE1<br>*Cr*APE1-Cu |
| ***Cr*APE1$_{WT}$** | 1:10 | | 17345<br>17407 | apo *Cr*APE1<br>*Cr*APE1-Cu |
| ***Cr*APE1$_{H163A}$** | - | 17280 | 17279 | apo *Cr*APE1 |
| ***Cr*APE1$_{H163A}$** | 1:1 | | 17279 | apo *Cr*APE1 |
| ***Cr*APE1$_{H163A}$** | 1:4 | | 17279 | apo *Cr*APE1 |
| ***Cr*APE1$_{H163A}$** | 1:10 | | 17277<br>17339 | apo *Cr*APE1<br>*Cr*APE1-Cu |
| ***Cr*APE1$_{C128S}$** | - | 17330 | 17329 | apo *Cr*APE1 |
| ***Cr*APE1$_{C128S}$** | 1:1 | | 17329<br>17390 | apo *Cr*APE1<br>*Cr*APE1-Cu |
| ***Cr*APE1$_{C128S}$** | 1:4 | | 17279<br>17391 | apo *Cr*APE1<br>*Cr*APE1-Cu |
| ***Cr*APE1$_{C128S}$** | 1:10 | | 17277<br>17391 | apo *Cr*APE1<br>*Cr*APE1-Cu |

**Supplementary Table 2:** Mass of Cr APE1 proteins without treatment or with addition of different ratio of Cu(II). Obtained by Native MS, complement to Supplemental Fig. 12.

**Table S3.**

| Protein | Oligomeric state | % | Measured Mass (Da) |
|---|---|---|---|
| ***Cr*APE1$_{WT}$-Cu** | Monomer | 94 ± 1 | 17890 ± 367 |
| | Dimer | | 42080 ± 1545 |
| | Oligomer | | 164133 ± 15200 |
| ***Cr*APE1$_{C128S}$-Cu** | Monomer | 58 ± 4 | 17677 ± 681 |
| | Dimer | | 37563 ± 1342 |

**Supplemental Table 3.** MALS analysis of oligomeric forms of CrAPE1 with and without addition of Cu for APE1$_{WT}$ and APE1$_{C128S}$

**Table S4.**

| Fraction | Name | Conc. | Cu mg/L | Cu µM | Molar ratio Cu:protein |
|---|---|---|---|---|---|
| Post-dialysis | complex *Cr*APE1$_{WT}$ - CuPCC1 | 150 µM | 82487.74 (+/-243.14) | 130 µM | 0.87 |
| F5 | *Cr*APE1 | 2 µM | 1066.91 (+/-9.44) | 1.68 µM | 0.84 |
| D11 | PCC1 | 57 µM | 21335.68 (+/- 119.88) | 33.59 µM | 0.59 |

**Supp Table 4.** ICP data for APE1-PCC1 complex pre and post cation-exchange column separation.

**Table S5.**

| **Primer number** | **Target** | **Sequence (5’-3’)** |
|---|---|---|
| P1 | *Cre16.g665250* (*ape1*) | AGCTCTTCTTGCTCGCTCCT |
| P2 | *Cre16.g665250* (*ape1*) | TACCGGCGTGCATTCCTATC |
| P3 | *Cre05.g248600* (*pcc1*) | GGGCTGGGAATTCGGTTCTG |
| P4 | *Cre05.g248600* (*pcc1*) | AGGGAGGACTGACGGTAGAG |
| P5 | *Cre10.g424775* (*ctp2*) | GGGTTGGGCTTTATTGGTGG |
| P6 | *Cre10.g424775* (*ctp2*) | CCATGTTGCGGACCTCCC |
| P7 | *Cre10.g422201* (*ctp4*) | CGTGTCTCCTGTTGCATACG |
| P8 | *Cre10.g422201* (*ctp4*) | GTGGAGGGATGGCAGTTCAT |
| P9 | *CIB1* cassette | CAGGCCATGTGAGAGTTTGCC |
| P10 | *CIB1* cassette | GCACCAATCATGTCAAGCCT |

**Supplemental Table 5.** PCR Primers used in this study

**Supplementary Data Sets (separate files)**

1. Data S1. *APE1* Co-expression data in 3 species.
2. Data S2. Superoxide detoxification assays

**Supplementary References also included in main references**

38. Harris EH, Stern DB, Witman G, Harris EH, Harris EH. The Chlamydomonas sourcebook. 2nd ed. Amsterdam Boston: Academic Press; 2009. 3 p.
